# Surrogate variables and the risk of overfitting in neuropsychiatric gene expression analyses

**DOI:** 10.64898/2026.10.05.756826

**Authors:** Dung T. Hoang, Ryan C. Thompson, Anina N. Lund, Eli Kritzer, Ariela S. Buxbaum-Grice, Divya John, Girish N. Nadkarni, Negar Golestani, Alexander W. Charney, Noam D. Beckmann

## Abstract

The reproducibility crisis in biomedical research calls for re-evaluation of how common analytic approaches affect results. One such approach, surrogate variable analysis (SVA), is widely used to adjust for unmeasured drivers of variation in transcriptomic differential expression analyses. We systematically quantified the impact of SVA using synthetic data with known ground truth and fourteen brain RNA-seq and microarray datasets spanning Alzheimer’s disease, autism spectrum disorder, bipolar disorder, schizophrenia, and Parkinson’s disease. In synthetic data, SVA usage frequently led to high false positive rates. This issue was not revealed by standard diagnostics nor resolved by strategies such as maximizing cross-dataset signal correlation. In real data, using SVA reduced cross-cohort reproducibility of differential expression signatures regardless of the number of surrogate variables included. These findings show that SVA risks inducing overfitting at realistic signal levels, advocating for a cautious, case-by-case application, with a preference for modeling known covariates.

## Introduction

Biomedical research faces a reproducibility crisis, with widespread failures to replicate published findings^1–3^, including in genomic studies, where methodological variations have led to conflicting results^4^. Differential expression analysis (DEA) has been extensively used in brain gene expression studies to identify differentially expressed genes (DEGs) associated with major neuropsychiatric disorders including Alzheimer’s disease (AD)^5–7^, autism spectrum disorder (ASD)^8, 9^, bipolar disorder (BP)^10–12^, Parkinson’s disease (PD)^13–15^, and schizophrenia (SZ)^16–18^. Although these studies have generated valuable insights into disease^5, 6, 10, 12, 14, 15, 19^ and informed therapeutic development^19–21^, attempts to reproduce findings in independent datasets often yield inconsistent results. This lack of reproducibility hinders the translation of findings into clinical applications and highlights the urgent need to characterize popular methods that may compromise reproducibility.

Identifying DEGs remains challenging because gene expression datasets are often confounded by unwanted biological and technical variation, such as environmental influences and batch effects^22–24^, that, if unaccounted for, can obscure true signals, reduce statistical power, and lead to misleading conclusions^25, 26^. Two broad strategies are used to account for this unwanted variation^27^. The first explicitly adjusts for known, measured covariates (e.g., batch, age, sex) using regression frameworks^27, 28^. While conceptually straightforward, this approach can be analytically involved and time-intensive and may miss important unmeasured variables^29^. The second strategy infers unmeasured sources of variation from the data’s covariance structure (i.e., hidden factors)^30–34^. Methods in this category include surrogate variable analysis (SVA)^35–37^, probabilistic estimation of expression residuals (PEER)^38^, and remove unwanted variation (RUV)^32, 39–41^. While these methods do not require manual covariate selection, they can compromise biological interpretability and statistical accuracy^29^, and recent studies show that they can introduce bias and false positives in gene expression analyses^42, 43^. Thus, it is essential to understand the trade-offs of inferring and adjusting for hidden factors in DEA.

Due to its ease of use and computational speed, SVA is among the most widely used methods for identifying hidden factors in gene expression studies^44–54^. SVA estimates both the number and values of latent variables (referred to as surrogate variables [SVs]) from gene expression data, which are then included as covariates in downstream DEA^35, 36^. Although SVA was shown to increase statistical power in small synthetic datasets (~1,000 genes, 20 samples,

≤2 SVs; here, “synthetic data” denotes computer-generated expression matrices with known ground truth and controlled noise/effect sizes) and in microarray datasets^35, 43^, its impact on DEA in larger, more complex datasets remains to be established^44–54^. SVA learns SVs from residual variation after removing the effect of the phenotype of interest, while permitting SVs to retain correlation with that phenotype to mitigate overfitting^35–37^. Using these SVs estimated from the data in the subsequent modeling of the same data to test for DEGs creates a circular dependency that can lead to overfitting (i.e., inflate false-positive rates [FPRs])^55^, but the specific circumstances under which such overfitting occurs are generally unknown. Thus, there is an urgent need to evaluate the impact of using SVA in DEA to ensure reliable and reproducible DEG results.

Here, we systematically evaluate the impact of using SVA in DEA using synthetic datasets and real transcriptomic datasets (RNA-seq and microarray) from fourteen brain cohorts spanning five major neuropsychiatric disorders (***Fig. 1***). Across synthetic data with controlled parameters, DEA with SVs yielded inconsistent, and often unacceptably high, FPRs relative to the conventional significance threshold of 0.05. In real datasets, where true DEGs are unknown, we used cross-dataset reproducibility as a validation metric and found that applying SVA consistently reduced reproducibility compared to analyses without SVA across most disorders. This pattern persisted when comparing DEGs identified in DEA with SVs to published DEGs results from independent datasets for the same disorders. Notably, these failures were not readily identifiable by inspecting p-value distributions, nor were they addressed by multiple-testing correction. Together, these findings indicate that SVA usage compromises reproducibility by unpredictably yet often inducing overfitting in DEA, and underscore the need for caution when using SVA in DEA.

**Fig. 1:**
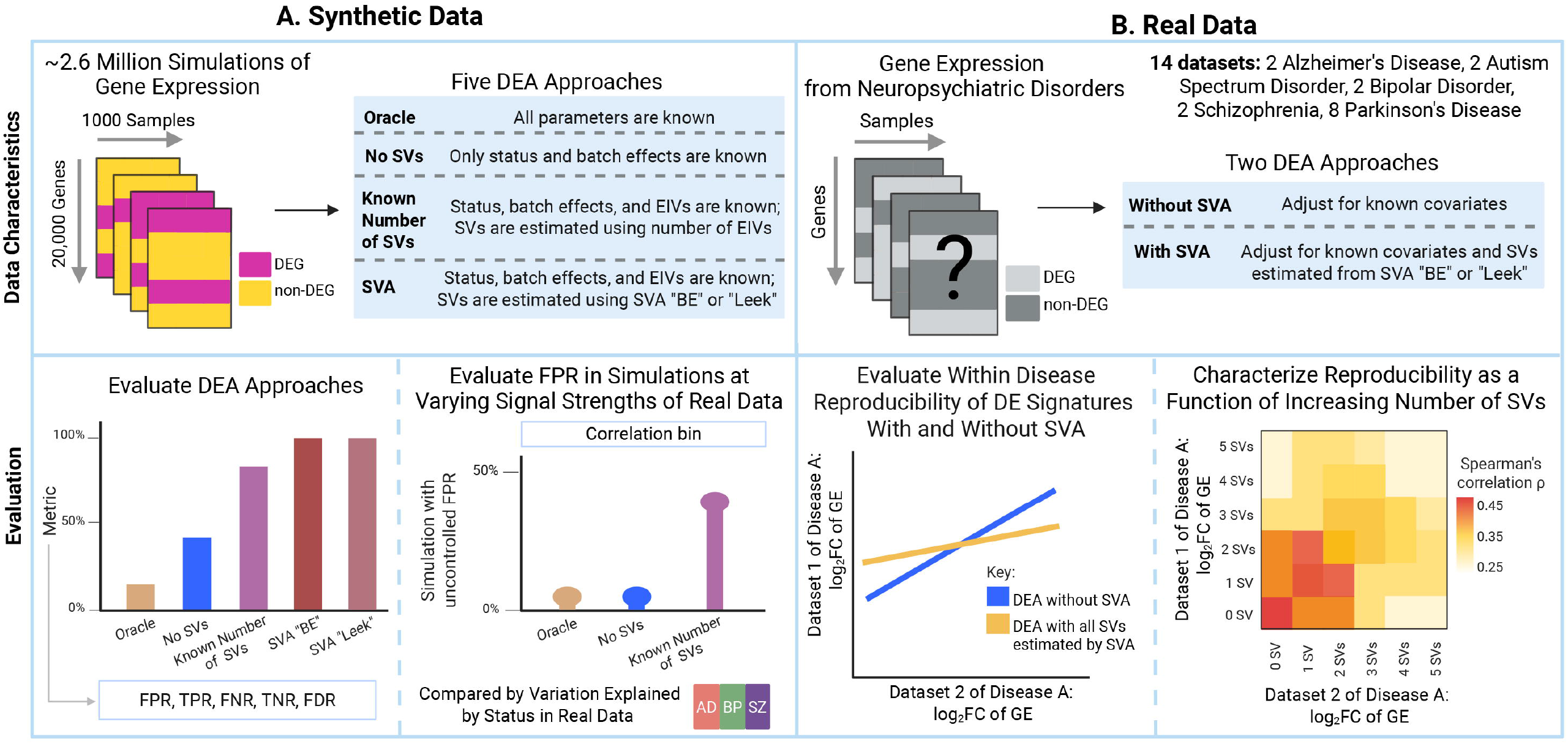
Study overview. The impact of surrogate variable analysis (SVA) on differential expression analysis (DEA) using both synthetic and real data. (**A**) **Synthetic data:** ~2.6 million gene expression simulations were generated, each containing 20,000 genes across 1,000 samples (500 cases, 500 controls) with known ground truth for differentially expressed genes. Five DEA approaches (defined in table) were applied that differ in their use of surrogate variables (SVs) and expression-influencing variables (EIVs), which are independent sources of hidden systematic variation that SVA aims to model. **Evaluation metrics:** false positive rate (FPR), true positive rate (TPR), false negative rate (FNR), true negative rate (TNR), and false discovery rate (FDR) were calculated. To connect synthetic data findings to real data, synthetic data performance was evaluated across signal strengths matching those observed in real datasets, defined by variance explained by disease status in Alzheimer’s disease (AD), bipolar disorder (BP), and schizophrenia (SZ). (**B**) **Real data:** Fourteen gene expression datasets were analyzed across five neuropsychiatric disorders where true DEGs are unknown (denoted by question mark) using two DEA approaches (defined in table). **Evaluation metrics:** DE signatures reproducibility was assessed by computing the Spearman correlations of log_2_fold-changes between datasets per disease, comparing results without SVA adjustment versus with SVA adjustment (using either all estimated SVs or incrementally increasing numbers of SVs).

## Results

### SV Inclusion in DEA Inflates FDR and FPR Beyond Acceptable Thresholds in Synthetic Data

Because real datasets lack ground-truth DEGs, we evaluated the effect of including SVs in DEA using synthetic gene expression data with known truth. We generated datasets with 20,000 genes for 1,000 individuals (500 cases, 500 controls), spanning varying signal-to-noise ratios and, among other varying parameters, expression-influencing variables (EIVs) representing unmeasured factors that induce unwanted variation in the data, yielding N=2,637,500 synthetic datasets (***Supplementary Table 1***). Each simulation constituted one dataset. Using variancePartition^56^, we confirmed that the range of variance explained by simulated parameters encompassed the values observed in real datasets (***Fig. 2, Extended Data Fig. 1, Supplementary Table 1***).

**Fig. 2:**
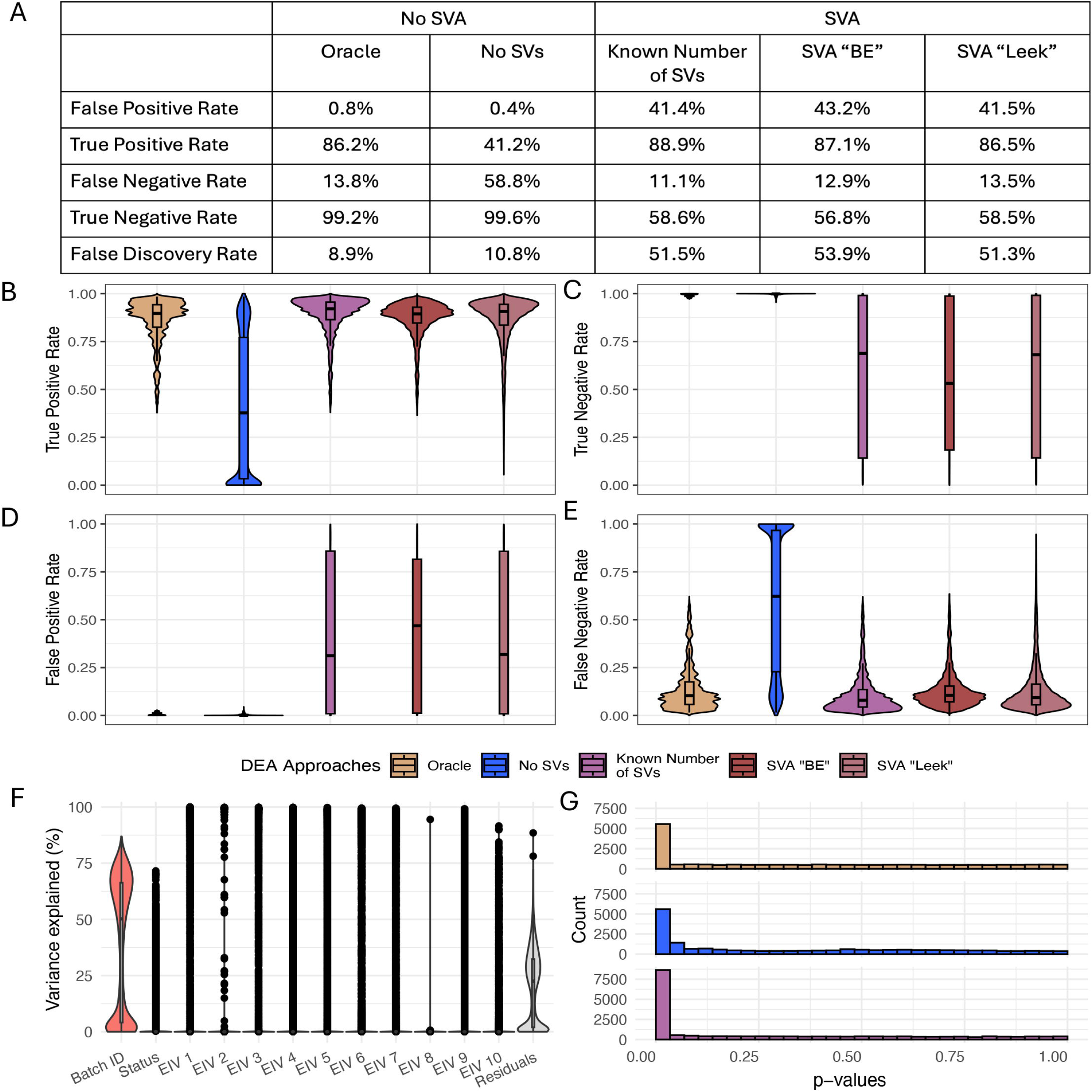
Methods using SVs yield high TPR and FPR. (**A**) Table summarizing the mean FPR, TPR, TNR, FNR, and FDR across five DEA approaches. (**B-E**) Violin and boxplots showing the distributions of TPR, TNR, FPR, and FNR (y-axis), respectively, for each method (x-axis) using violin boxplots. The color is defined by the legend. (**F**) The variance decomposition for batch, status, 10 EIVs, and residuals for one representative simulation. This simulation has a FPR of 0.235, and the following parameters: number of EIVs = 10; total SV variability = 2.5; variability per SV = 0.79; fraction of differentially expressed genes = 0.04; status effect size = 0.2; residual standard deviation = 0.3; fraction of genes affected per SV = 0.7; random seed = 12345. (**G**) Histogram showing the distribution of p-values for the same simulation shown in panel F, comparing three DEA approaches: Oracle, No SV, and SVA Known Number of SVs. Despite differences in modeling approaches, all three DEA approaches yield similar p-value distributions with uniform patterns and peaks near zero. The x-axis shows the p-values; the y-axis shows the count of the simulation. The color is defined by the legend.

To evaluate how incorporating SVs in DEA affects the identification of true DEGs, we compared five DEA modeling approaches spanning common covariate-adjustment strategies (***Fig. 1A***, see *Methods*): (1) an “Oracle” model adjusting for batch and all EIVs as they are known exactly; (2) a “No SVs” model adjusting for batch only; (3) “SVA Known Number of SVs,” adjusting for batch and SVs estimated from the data using the true number of EIVs; (4, 5) “SVA BE” and “SVA Leek,” adjusting for batch and SVs with the number of SVs chosen by the default “BE” method^57^ and the “Leek” method^58^, respectively. Because approaches 3-5 differ only in how the number of SVs is selected, results for “SVA BE” and “SVA Leek” are reported in the ***Supplementary Material*** unless otherwise specified. The “Oracle” and “No SVs” strategies serve as reference points for, respectively, the theoretically optimal performance if all EIVs are known, and the performance obtained without any attempt to model the EIVs. Performance of each model was evaluated using false discovery rate (FDR), FPR, true positive rate (TPR), false negative rate (FNR), and true negative rate (TNR, ***Fig. 1A***). Given that an increase in FDR in the context of overfitting is often caused by an increase in FPR, our study mainly focuses on FPR. We validated that our synthetic data generation approach was appropriate for characterizing SVA usage in DEA by showing that it can recapitulate the results presented in the original SVA manuscript when using comparable parameters^59^ (***Supplementary Material*** *“Our synthetic data framework reproduces original SVA results”*).

To establish expectations for DEG identification, the “Oracle” and “No SVs” approaches were first characterized. As expected, the “Oracle” approach outperformed all others, yielding a high TPR while controlling FPR (mean FPR = 0.8%, TPR = 86.2%, FNR = 13.8%, TNR = 99.2%, FDR = 8.9%, ***Fig. 2A, B, C, D, E, Supplementary Table 1***). The “No SVs” approach also controlled FPR but showed the widest TPR range (mean FPR = 0.40%, TPR = 41.2%, FNR = 58.8%, TNR = 99.6%, FDR = 10.8%, ***Fig. 2A, B, C, D, E, Supplementary Table 1***). In contrast, SVs adjusted approaches increased TPR but failed to control FDR and FPR on average (“SVA Known Number of SVs”: mean FPR = 41.4%, TPR = 88.9%, FNR = 11.1%, TNR = 58.6%, FDR = 51.5%, ***Fig. 2A, B, C, D, E, Supplementary Table 1***). This poor FDR and FPR control are not explained by inaccurate estimation of the number of SVs, as SVA generally estimates the true number of simulated EIVs accurately (***Extended Data Fig. 1, Supplementary Material*** *“SVA generally estimates the number of simulated EIVs accurately”*). Nevertheless, 24.08% and 39.11% of simulations using SVA approaches maintained FDR and FPR below the conventional 5% threshold. Consistent with these patterns, Oracle and No SVs maintained high TNR, whereas “SVA Known Number of SVs” achieved a low FNR at the expense of TNR (***Fig. 2A, C, E***). These results indicate that the apparent TPR gains with SV adjustment represent a false improvement as higher sensitivity is achieved by overfitting to noise, as demonstrated by its inability to control FDR and FPR within acceptable statistical thresholds.

To assess whether DEA behaves as expected, researchers often inspect p-value distributions (uniform under the null, and with an excess of small p-values when true DEGs are present) and quantify deviations from uniformity^35, 60, 61^. We examined p-value distributions for five randomly selected simulations with 10 EIVs (***Fig. 2F, G, Extended Data Figure 2***). Across all approaches, deviations from uniformity were similar and consistent with the true signal, regardless of whether overfitting occurred. This emphasizes that p-value distributions alone are insufficient to detect overfitting when SVs are used in DEA, since true signals and false positives from overfitting result in distributions with similar shapes.

### High FDRs Persist at Signal-to-Noise Ratios Characteristic of Real Data

Given that an increase in FPR causes an increase in FDR, we examined FDR patterns to assess whether high FPR and thus high FDR persisted under signal-to-noise ratios characteristic of real data. Because loss of FDR control in DEA with SVA may depend on the strength of the true differential expression (DE) signal, we evaluated DEA performance across synthetic datasets grouped into deciles based on signal strength (defined as the percentage of total variance explained by the phenotype) and quantified TPR, FPR, and FDR for each group (***Fig. 3***). Distinct patterns emerged across the “Oracle,” “No SVs,” and “SVA Known Number of SVs” models as signal increased (***Supplementary Material, Extended Data Fig. 3)***. As expected, “Oracle” consistently maintained low FPR, low FDR, and high TPR across all signal strength deciles. “No SVs” also maintained low FPR and FDR but at the cost of a lower TPR at lower signal strengths (***Supplementary Material, Extended Data Fig. 3)***. In contrast, “SVA Known Number of SVs” maintained high TPR in 98.79% of the simulations (TPR ≥ 0.5; TPR range: 96.74%-100%), but frequently lost type I error control: 59.64% had FPR > 0.05; FPR (range: 3.89%-92.48%), and 75.65% had FDR > 0.05 (FDR range: 10.36%-99.19%, ***Fig. 3***). Although stronger signal mitigated this trade-off, elevated FDRs and FPRs remained common across much of the parameter space.

**Fig. 3:**
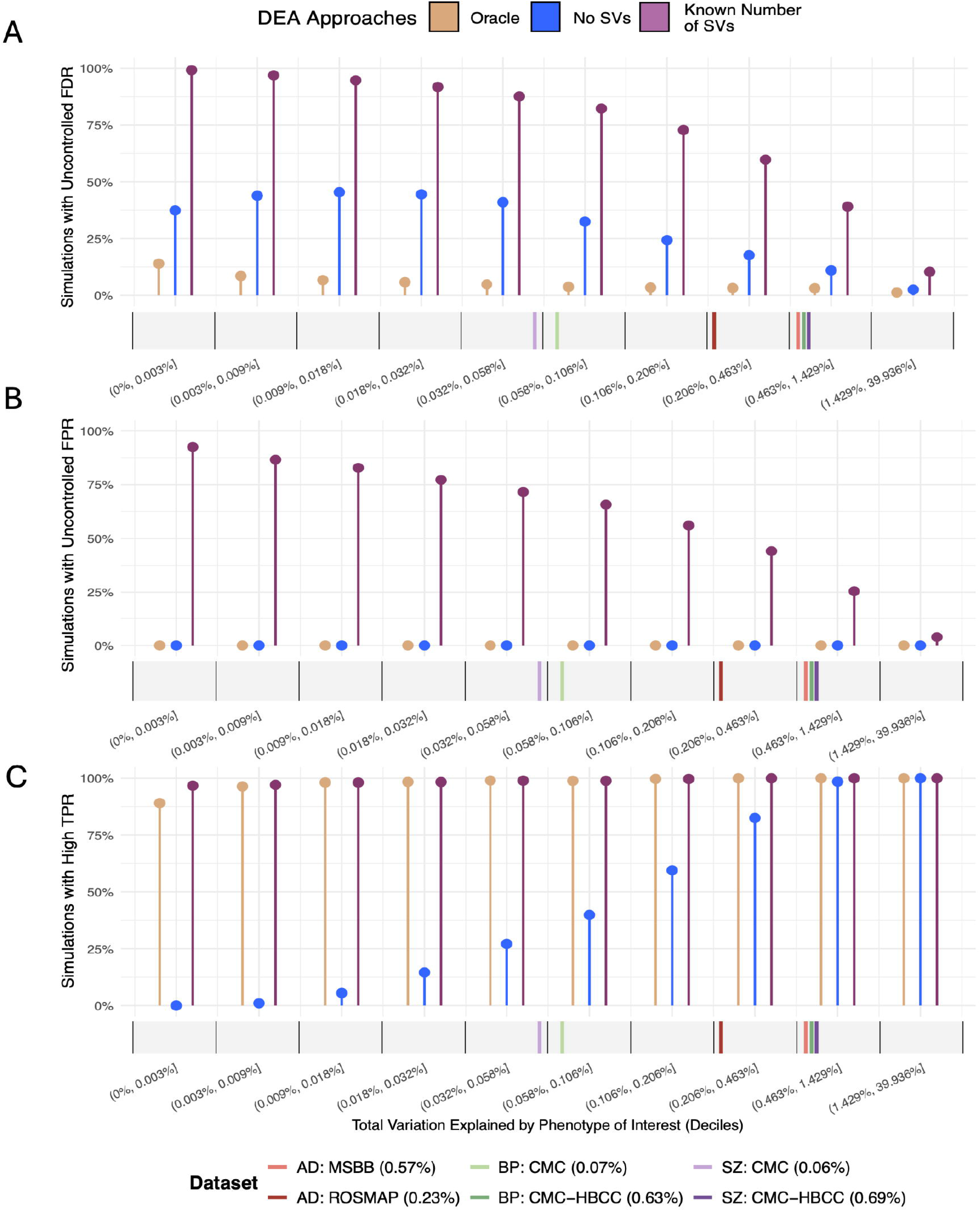
There are a high number of simulations with uncontrolled FPR and FDR at real signal strengths. Lollipop plots showing the proportion of simulations meeting three criteria: (**A**) uncontrolled FDR (i.e., FDR < 0.05, 38% of total simulations), (**B**) uncontrolled FPR (i.e., FPR < 0.05, 20% of total simulations), and (**C**) high TPR (i.e., TPR ≥ 0.5, 72% of total simulations) across three DEA approaches (defined in the legend). Synthetic data are stratified by signal strength, defined as the total variance explained by the phenotype of interest (i.e. disease status) and grouped into deciles. The x-axis is the percent variance explained by the phenotype of interest within each decile; the y-axis is the simulations meeting each criterion. For comparison, six real datasets from Alzheimer’s disease (AD), bipolar disorder (BP), and schizophrenia (SZ) studies are overlaid at their observed variance explained by disease status (values shown in parentheses). The datasets are: **AD:** Religious Orders Study and Rush Memory and Aging Project (ROSMAP) and Mount Sinai/JJ Peters VA Medical Center Brain Bank (MSBB); **BP and SZ:** Common Mind Consortium datasets from Mount Sinai, University of Pennsylvania, and University of Pittsburgh Brain Banks (CMC) and Common Mind Consortium from the National Institute of Mental Health’s Human Brain Collection Core (CMC-HBCC).

To set biologically relevant signal strengths, we benchmarked our simulation range against real neuropsychiatric datasets, defining signal-to-noise as the median percent variance explained by the phenotype across genes. We used four large RNA-seq cohorts: 1) Religious Orders Study and Memory and Aging Project (ROSMAP) data^62, 63^; 2) Mount Sinai Brain Bank (MSBB) data^64^; 3) CommonMind Consortium Mount Sinai/University of Pennsylvania/University of Pittsburg (CMC) data^65^; 4) CommonMind Consortium National Institute of Mental Health’s Human Brain Collection Core (CMC-HBCC) data^65^. The observed percents variance explained were 0.23% (ROSMAP) and 0.57% (MSBB) for AD; 0.07% (CMC) and 0.63% (CMC-HBCC) for BP; 0.06% (CMC) and 0.69% (CMC-HBCC) for SZ (***Fig. 3***). To ensure comprehensive signal strength coverage, we mapped these values to deciles of our simulation distribution and set the assessment window to the full interval spanning those deciles (0.032%-1.429%). Within this biologically meaningful range, the “SVA Known Number of SVs” approach achieved high TPR (range: 98.96%-100%, ***Fig. 3C***) but retained unacceptably high error rates (FPR ranges: 25.46%-71.56%; FDR ranges: 39.07%-87.63%, ***Fig. 3A, B***). These results demonstrate that the poor FPR control with SVA in our DEA of synthetic data is not an artifact of implausible simulation parameters; it persists at signal strengths characteristic of real neuropsychiatric data.

### Maximizing Disease Signal Correlation Does Not Improve FPR Control in Synthetic Data

Maximizing cross-dataset DE signatures correlation has been proposed as a strategy to select the optimal number of SVs for SVA-based DEA and to improve reproducibility^44^. To assess whether this approach effectively controls FPR, we simulated 500 pairs of independent datasets with a shared true signal but unrelated noise structures (hereafter referring to the two data sets in a given pair as Simulated Data 1 and Simulated Data 2; see *Methods*, ***Fig. 4, Extended Data Fig. 4***). DEA was performed using SVA with number of SVs ranging from 0 to the true number of simulated EIVs (N = 25), and log_2_fold-change (log_2_FC) concordance between the DE signatures from the two datasets evaluated using Spearman correlation. The mean percent variance explained spanned 0.027% to 0.667%, corresponding to signal strengths simulated with parameters ranging from 0.1 to 0.5, respectively.

**Fig. 4:**
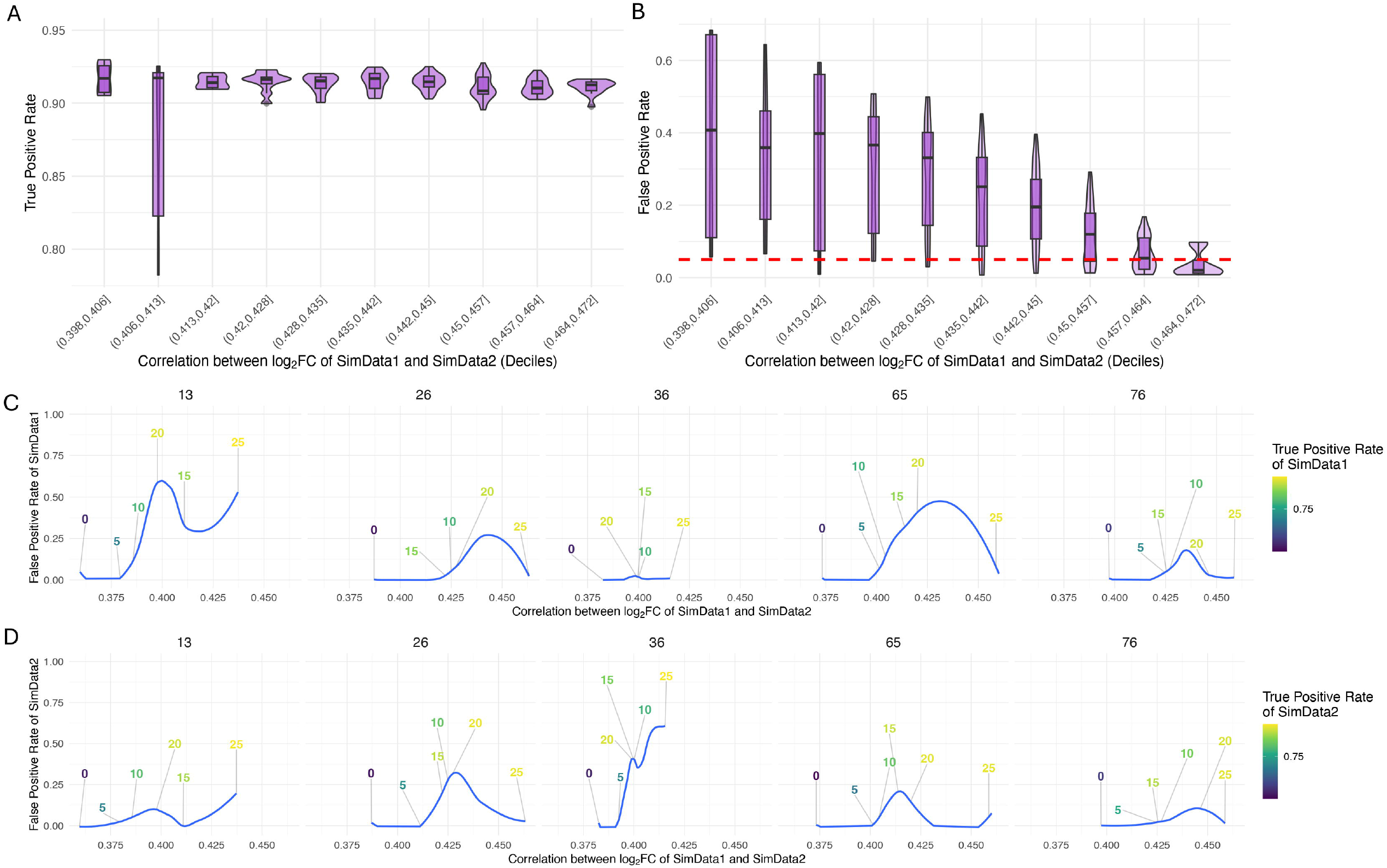
Higher correlation of log_2_FC between independent simulated datasets does not guarantee FPR control. Pairs of simulated datasets (termed SimData1 and SimData2) were generated with identical signal and number of true DEGs but different noise structures. (**A**-**B**) Violin and boxplots showing the distribution of the correlation of log_2_FC between SimData1 and SimData2 in deciles (x-axis) against either (**A**) TPR across all simulated data or (**B**) FPR across all simulated data (y-axis). The red dashed line in (**B**) indicates FPR = 0.05. (**C**-**D**) Line plots for five randomly selected simulation seeds (the top number of each plot) illustrate that FPR patterns vary unpredictably across seeds, even when correlations remain high. LOESS curves (blue) show the FPR (y-axis) as a function of log_2_FC correlation (x-axis), with numbers along each curve indicating the estimated number of SVs (0–25), and numbers are colored according to their TPR values (defined in the legend). The true number of simulated EIVs was 25, with the mean percent variance explained = 0.241% (disease status effect size of 0.3; See *Methods*). This effect size corresponds to a mean percent of total variance explained of 0.24% for the phenotype of interest, aligning with the signal strength observed in real datasets.

At any fixed signal strength, increasing number of SVs used in DEA resulted in higher log_2_FC correlations between Simulated Data 1 and Simulated Data 2 (***Extended Data Fig. 4A***). However, this apparent gain in reproducibility did not corresponds to higher TPR at all signal strengths (***Extended Data Fig. 4B***). Instead, TPR was only higher when the underlying signal increased (***Extended Data Fig. 4B***). Particularly, this relationship was observed at signal strength comparable to that observed in real data such as ROSMAP (synthetic data signal strength: 0.3 with mean percent variance explained = 0.241%; ROSMAP signal strength: 0.23%) (***Fig. 4A***). Although not associated with a better TPR, an increase in log_2_FC correlations was observed with a decreased in FPR, with the lowest FPR values at the maximum correlation (***Fig. 4B, Extended Data Fig. 4C***). However, even in the highest correlation bin (0.464-0.484], 25% of simulations still exceeded the 0.05 FPR threshold (***Fig. 4B***), indicating that maximizing cross-dataset correlation in DEA results obtained while using SVA both does not reliably control FPR and incorrectly implies improved performance.

To investigate relationships between FPR, TPR, log_2_FC correlation, and number of SV, five randomly selected simulation instances for each of the two simulated datasets were inspected (***Fig. 4C, D***). No stable or monotonic relationships emerged among these metrics (***Fig. 4C, D***), consistent with overfitting. These findings show that although approaches maximizing cross-dataset log_2_FC correlations can lower FPR without improving TPR, they fail to ensure acceptable control for false positives in a substantial proportion of analyses.

### SVA use in DEA Reduces DEG Reproducibility across Neuropsychiatric Disorders

To determine whether using SVA for DEA can induce overfitting in real data, we analyzed fourteen public brain gene expression cohorts across five neuropsychiatric disorders (RNA-seq and microarray data with and without SVA^44, 48, 65–71^. We compared (1) a baseline model that controls for the same covariates as each original study^44, 48, 65–71^ (***Supplementary Table 2***) and (2) an SVA-adjusted model including SVs estimated by the “BE” or “Leek” procedures to the baseline model (***Supplementary Table 2***). Because true DEGs are unknown, direct assessment of TPR and FPR is not possible. Instead, we quantified the reproducibility of DE signatures non-parametrically using Spearman’s ρ between log_2_FC values from independent brain gene expression datasets for the same disorder (two RNA-seq datasets for each of AD, ASD, BP, and SZ, and eight microarray datasets for PD), noting that higher reproducibility reflects both higher TPR and lower FPR^72^. To assess the effects of phenotype and published covariates in each dataset, we partitioned the variance in the gene expression data attributable to each factor (***Extended Data Fig. 5***).

Overall, SVA reduced reproducibility in four of five disorders (AD, BP, PD, and SZ, ***Fig. 5***). For AD, the trait with the highest reproducibility in the baseline model, cross-dataset correlation dropped from ρ = 0.46 (baseline) to ρ = 0.12 (with SVA), and BP and SZ showed similar declines (***Fig. 5A***). This drop persisted when using an alternative of SVA designed to ensure convergence of the SV estimation in AD datasets^73^ (***Supplementary Material*** *“Convergence-based SV estimation reduces reproducibility of AD DE signatures,”* ***Extended Data Fig. 8***). ASD was the exception: DE signatures failed to replicate with or without SVA (***Fig. 5A***). For PD, albeit smaller, lower reproducibility was still generally observed when using SVA (range of ρ_withoutSVA_ - ρ_withSVA_ = −0.03 to 0.11) (***Fig. 5B***). Due to sample size considerations, subsequent analyses were performed on only the largest datasets in which reproducibility was observed in the baseline models not using SVA: SZ, AD, and BP.

**Fig. 5:**
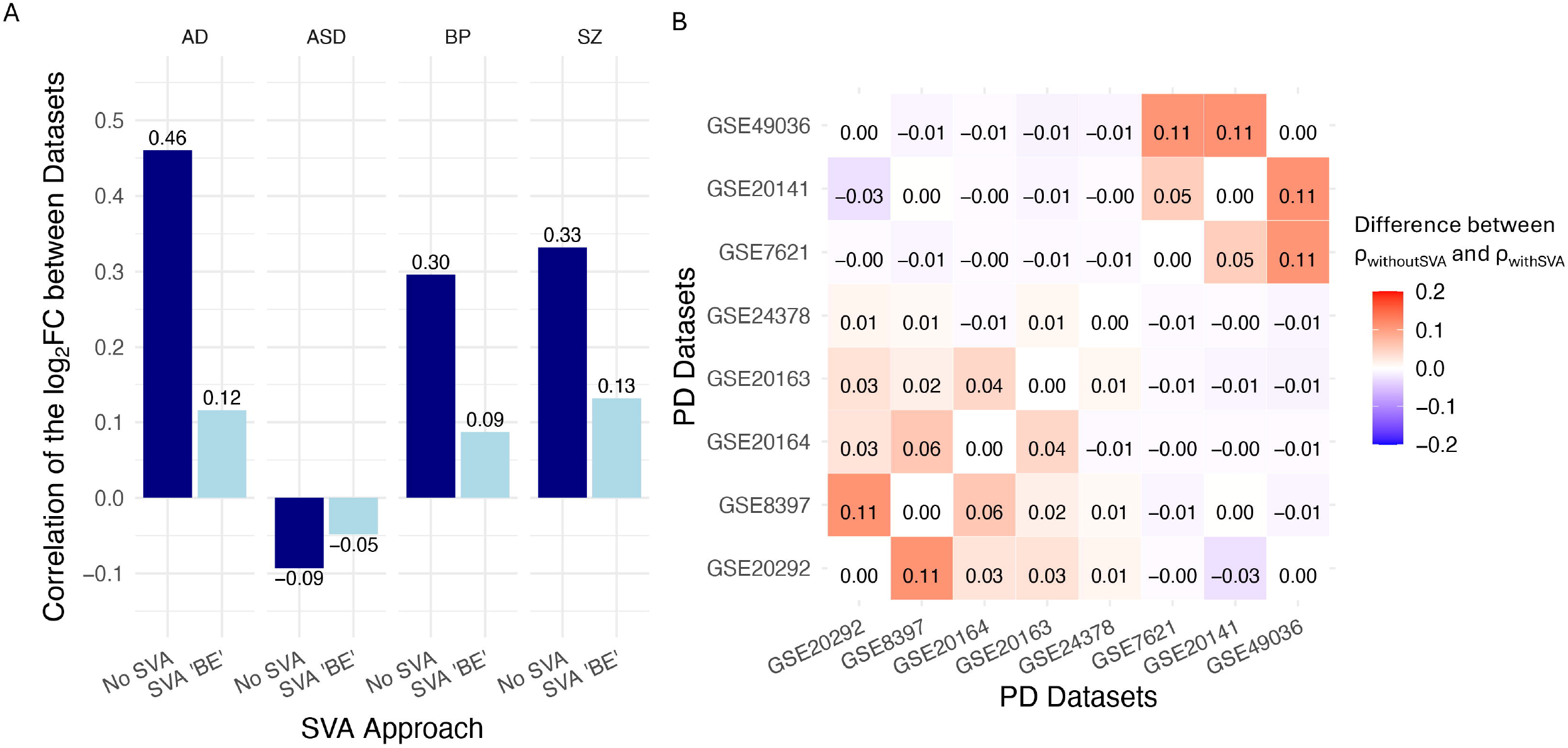
DEA with SVA Reduces Cross-Datasets Reproducibility in Neuropsychiatric Disorders. (**A**) Bar plots showing the Spearman correlations of log_2_FC (x-axis) between independent dataset pairs for Alzheimer’s disease (AD), autism spectrum disorder (ASD), and bipolar disorder (BP) with SVA “BE” adjustment (light blue) and without SVA adjustments (navy) (y-axis). Numbers on top or below the bars indicate the actual Spearman ρ. Lower Spearman ρ with SVA adjustment indicates reduced reproducibility of DE signatures between datasets. (**B**) Heatmap displays the change in log_2_FC correlations (with SVA minus without SVA) for all pairwise comparisons across eight independent Parkinson’s disease microarray datasets (x-axis and y-axis). Negative values (blue) indicate that SVA reduced correlation between dataset pairs, while positive values (red) indicate increased correlation. Two clusters are observed in this analysis: **Cluster 1**: GSE49036, GSE20141, and GSE7621; **Cluster 2**: GSE24378, GSE20163, GSE20164, GSE8397, and GSE20292. Each cluster contained results from datasets generated by a different microarray platform (GPL570 for **Cluster 1** and GPL96 and GPL1352 for **Cluster 2**), emphasizing differences in reproducibility for the same disease between measurement platforms.

In all cases, the SVs inferred by SVA were approximately orthogonal to the phenotypes - an undesirable property, as such orthogonality prevents them from capturing true phenotype-associated confounders, leading to overfitting and reduced reproducibility. (***Supplementary Material*** *“Despite overfitting protection, SVs remain nearly orthogonal to phenotype,”* ***Extended Data Fig. 6A***). Consistent with overfitting occurring only when true signal is present, overfitting disappeared in ROSMAP when a null was generated by shuffling true disease labels (***Supplementary Material*** *“SVA does not lead to overfit under null conditions,”* ***Extended Data Fig. 6B***). Finally, we tested whether the reproducibility decline could be explained by overestimation of the number of SVs (the smallest “BE” selection across AD, BP, and SZ was 27 SVs in MSBB). Including any number of SVs in the model from 1 up to the maximum selected by SVA “BE” (e.g., 27 SVs in MSBB) reduced reproducibility (***Fig. 6A, Extended Data Fig. 7C-E***). These results indicate that the reduced reproducibility observed in these data sets is intrinsic to SV adjustment in DEA, consistent with overfitting, rather than an artifact of selecting too many SVs.

**Fig. 6:**
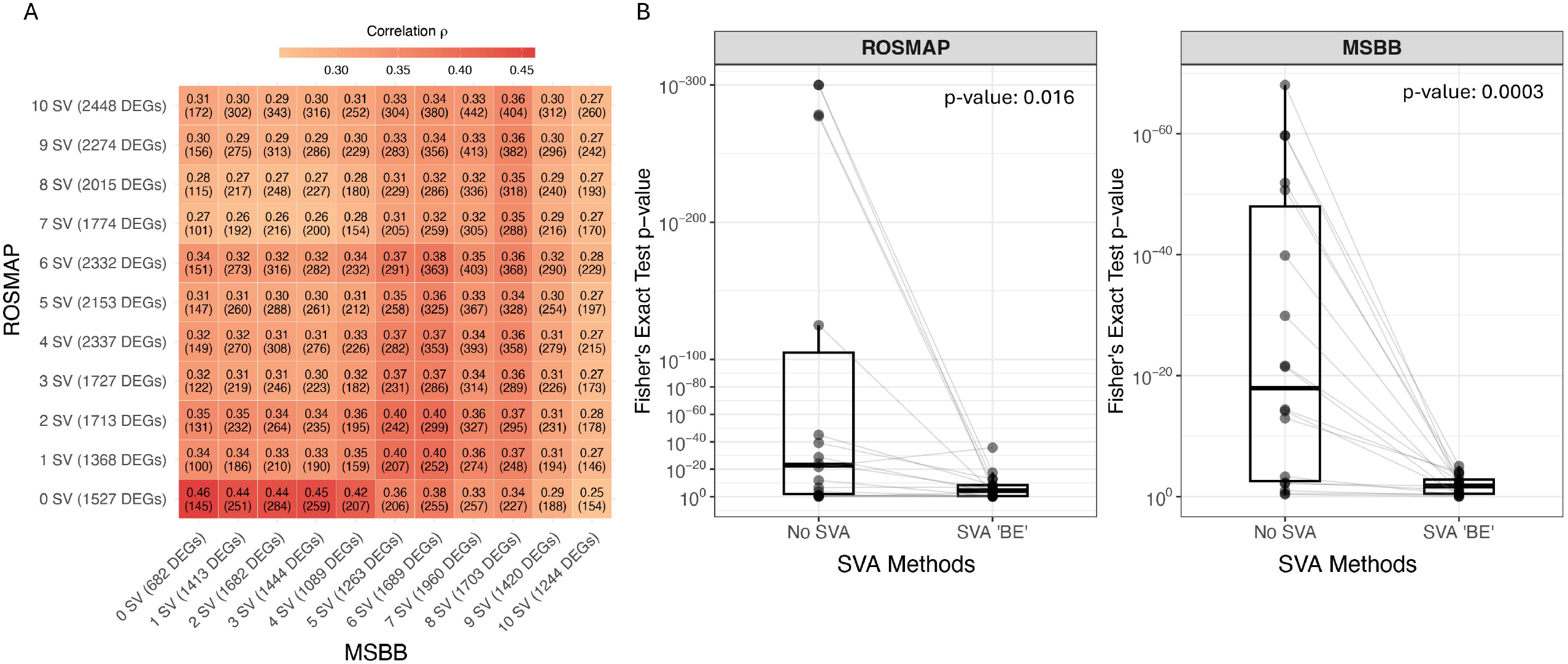
SVA Reduces Reproducibility of AD DE Signatures. (**A**) Heatmap showing Spearman correlations of log_2_FC between ROSMAP (x-axis) and MSBB (y-axis) when incrementally including SVs (0-10 SVs). Each cell displays the Spearman ρ values with the number of overlapping significant DEGs in parentheses. Higher Spearman ρ and more overlapping DEGs indicate better reproducibility between datasets. (**B**) Box plots comparing the statistical significance of DEGs overlap between the two primary AD datasets (ROSMAP and MSBB) and ten additional publicly available AD datasets. The x-axis shows two methods: No SVA (left) and SVA “BE” (right), where SVA “BE” estimated 27 SVs for MSBB and 48 SVs for ROSMAP; the y-axis shows p-values from Fisher’s exact tests (negative log scale), where smaller p-values indicate more significant overlap (better reproducibility). The p-values from Wilcoxon signed-rank tests are displayed in each facet.

To test whether the observed decline in reproducibility generalizes beyond the primary cohorts, we benchmarked DE signatures computed with and without SVA against independent published DEG lists (9 in AD^74–82^, 5 in BP^10, 83–86^, and 5 in SZ^87–90^, ***Supplementary Materials***). Enrichments were generally weak, consistent with the limited power in DEG discovery. However, in AD and SZ, comparisons of enrichment statistics revealed that using SVA significantly reduced overlap with previously published DEGs (***Fig. 6B, Supplementary Materials, Extended Data Fig. 7A, B***). These results confirm that incorporating SVA into DEA reduces reproducibility relative to analyses without SVA.

## Discussion

Our study investigates a widely used class of methods that infer unmeasured confounders directly from expression data, conditioned on the phenotype of interest, to enhance detection of disease-associated changes. Focusing on SVA, we systematically evaluated its impact on DEA in synthetic and real data and found that it frequently overfits (i.e., inflates FPR and decreases reproducibility). In synthetic data, this overfitting escaped detection *via* standard diagnostics (e.g., inspecting p-value distributions) and was not reliably mitigated by choosing SV numbers *via* cross-dataset DE-signal correlation. Both when using the true confounder number (“SVA Known Number of SVs”) and when estimating it via “BE” or “Leek,” the gain in sensitivity provided by SVA came at the cost of a recurrent loss of FPR control. Even in the best-case “SVA Known Number of SVs” setting, roughly two in five reported DEGs were false positives, and this poor FPR control persisted when simulations were calibrated to effect sizes typical of neuropsychiatric datasets, suggesting similar FPR inflation in real data. Consistent with this, including SVs in DEA models reduced cross-dataset reproducibility of DE signatures in fourteen RNA-seq and microarray datasets spanning five neuropsychiatric disorders. Moreover, reproducibility declined regardless of the number of SVs included.

Approximately 40% of simulations that incorporated SVs kept FPR below the 0.05 threshold. Despite extensive profiling, we did not find stable patterns that explained why overfitting was absent in this minority. This suggests that there may exist a specific, uncharacterized set of conditions under which SVA usage in DEA avoids overfitting. This aligns with our replication of the original SVA simulations, where FPR was well controlled under small-sample, few-SV settings that were representative of datasets generated at that time^59^. In contrast, modern RNA-seq studies typically involve larger cohorts and more complex designs, yielding many more estimated SVs^44–54^. While newer SVA-based methods, including for RNA-seq^37^, have since been proposed, they still estimate SVs using knowledge of the phenotype and thus remain vulnerable to overfitting^37, 91–93^. Our simulations also indicate that cross-dataset reproducibility of DE signatures can reduce the likelihood of overfitting when using SVA. However, even at the highest reproducibility observed in simulation (representing an upper bound for real data) FPR was often not controlled, so this strategy cannot guarantee absence of overfitting.

Analyses of real neuropsychiatric datasets further corroborated this pattern: including SVs in DEA models systematically reduces cross-study reproducibility of DE signatures. The decline appears both when using the algorithm’s recommended number of SVs and when including only a few SVs, indicating that the loss of reproducibility stems from SV inclusion itself rather than simply from using too many SVs. The consequences of this loss of cross-study reproducibility are substantial: more false discoveries, reduced generalizability to independent datasets, and impaired clinical translation. Practical implications are potentially severe: Poor reproducibility directly impedes biomarker development and therapeutic target validation, resulting in wasted resources pursuing false leads that appear promising in initial studies but fail during validation, ultimately misdirecting scientific effort and delaying genuine therapeutic advances that could otherwise contribute to more effective treatments for neuropsychiatric patients.

The overfitting observed when using SVA likely reflects a broader issue shared by methods that estimate hidden confounders while incorporating the phenotype of interest into the model used to infer those confounders^31^. This setup creates data leakage: information from the outcome (e.g., phenotype of interest) contaminates intermediate steps that should be independent, even when safeguards to limit overfitting are used^44^. Such leakage can inflate apparent model performance and elevate FPR, including in well-powered studies^2^. Similar concerns have been raised for SVA and related approaches^42, 73^, and newer methods have been proposed to address unwanted variation in large-scale RNA-seq data^60, 73, 91, 93^. Rigorous, head-to-head evaluation of these newer approaches will be essential to identify which remove unwanted variation without introducing overfitting.

Our study has several limitations. Although we ran ~2.6 million simulations, they cannot fully recapitulate the complexity of real-world datasets. This concern is partially mitigated by our simulation calibrations to match signal strengths and co-expression structure observed in real neuropsychiatric datasets. Future work will need to explore further simulated conditions that are not limited by some of the assumptions present here, including linear relationships between confounders and gene expression, normally distributed normalized gene expression values, and independence between confounding variables. To maintain model simplicity and interpretability, we only included one known covariate, batch, to be included in DEA with and without SVA. Although we did not test including additional known covariates when using SVA in synthetic data, the decrease in reproducibility observed in real neuropsychiatric DEAs suggest overfitting would persist. Our analyses focused on standard linear model approaches to DEA. Future work is necessary to generalize our results to other machine learning-based methods. Reproducibility of DE signatures was quantified using Spearman correlation of log_2_FC values between datasets with the same disease. Although this non-parametric approach is well supported for this purpose^94^, alternative approaches to measuring cross-study reproducibility such as pathway consistency or functional enrichment patterns were not performed. Future work using these alternative reproducibility metrics may provide complementary insights. While our real-data validation spans multiple disorders and technologies, it was limited to brain transcriptomes and neuropsychiatric conditions. Future work is needed to determine whether the patterns observed here extend to all traits and tissue types. Finally, although literature support exists that extends our conclusions to other methods that estimate unknown confounders from gene expression data with knowledge of the phenotype of interest^42, 43^, systematic evaluation of the impact on overfit of using these methods for DEA will be required.

In conclusion, the use of SVA in DEA requires careful consideration, as our results indicate that this can lead to substantial difficult-to-detect overfitting that compromises reproducibility in both real and synthetic data. While SVA aims to adjust for hidden confounders, its estimated covariates should not replace rigorous quality control or thoughtful experimental design, since known covariates provide a more reliable and interpretable basis for adjustment. The extent of SVA-induced overfitting likely varies across datasets and contexts, meaning its impact can range from negligible to severe, with direct consequences for biological interpretation. Thus, the validity of previous findings obtained with SVA may need to be reassessed on a case-by-case basis, particularly in studies where overfitting cannot be ruled out. Whenever feasible, modeling known sources of variation explicitly, as opposed to relying on SVs, is a more robust strategy. Finally, to ensure the reliability and reproducibility of DEA results, validation through independent replication or alternative analytical methods is strongly recommended to minimize method-specific biases and strengthen confidence in the resulting biological conclusions.

## Methods

### DEA Simulations

Synthetic gene expression datasets were simulated with multiple sources of variation to systematically evaluate the effect of using SVA in DEA under varying levels of known and unknown confounding. These datasets represented differential expression experiments where case-vs-control DEA were conducted in the presence of both a known batch effect and EIVs representing true but unknown sources of systematic variation in the data. In total, 2,637,500 unique datasets were generated.

#### Defining the simulation parameter grid

Normalized gene expression data was simulated with the following parameters:

1. Constant parameters:
  - Number of molecular features (genes): n_features = 20,000
  - Number of individuals: n_individuals = 1,000 (500 cases and 500 controls)
  - Number of samples per individual: n_samples_per_individual = 1
  - Number of batches (defined as a categorical variable): n_batches = 15
  - Batch signal strength: standard deviation (SD) of batch effects: batch_sd = 0.3.
2. Varying parameters:
  - Number of expression influencing variables (EIVs): n_eiv ∈ {0, 10, 20,…,100}.
  Proportion of genes that are differentially expressed between case and control groups: fraction of case vs. control DEGs: fraction_degs ∈ {0,0.02,0.04,0.06,0.08,0.1,0.2,0.3,0.4,0.5}.
  - Expected fraction of genes affected by each EIV (emulating coexpression): fraction_eiv ∈ {0.1,0.3,0.5,0.7,0.9}.
  - Magnitude of the differences in gene expression between case and controls: status_sd ∈ {0.1,0.2,0.3,0.4,0.5}.
  - Total strength of the EIV-associated variation added to the simulated data: total_eiv_sd ∈ {0,0.5,1,1.5,…,10}.
  - Strength of the variation associated with individual 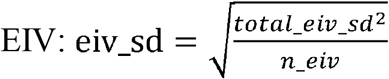. (If n_eiv is 0 then eiv_sd is not used.)
  - Strength of the random noise added to the simulated data: resid_sd ∈ {0.1,0.2,0.3,0.4,0.5}.

#### Generating synthetic datasets

For each combination of the above parameters, ten case-control datasets were simulated using different random seeds (sequential integers starting from 12345) through the following steps:

1. Generate IDs:
  a. Generate n_individuals distinct individual IDs.
  b. Generate n_batches distinct batch IDs.
  c. Generate n_eiv distinct EIV names.
2. Generate sample metadata:
  a. Assign one sample per subject.
  b. Randomly designate half of the subjects as cases and the other half as controls
  c. Generate n_eiv random EIV weights from exponential distribution *Exp*(1). Scale them so that their sum of squares equals total_eiv_sd^2^. Order them from largest to smallest and label them eiv_sd_1 through eiv_sd_n.
  d. Generate per-sample values for each EIV by drawing from *N* (0,eiv_sd_N). Repeat this for all EIVs using the corresponding scaled values eiv_sd_1 through eiv_sd_n.
  e. Assign each sample to a batch ranging from 1 to n_batch randomly and independently.
3. Generate feature metadata:
  a. Generate a list of features and marks the first n_features * fraction_degs as true DEG and all other features as non-DEGs.
4. Construct the feature data matrix:
  a. Batch effects: Form the batch design matrix as the one-hot encoding of the batch membership for each sample (n_samples x n_batches). Simulate a matrix of batch effects (n_batches x n_features) by drawing values from a *N* (0,batch_sd). Multiply the batch design matrix the batch effect matrix to obtain the batch contribution to expression.
  b. EIV effects: Form the EIV design matrix as a matrix (n_samples x n_eiv) of sample EIV weights. Simulate an EIV loading matrix (n_eiv x n_features) by drawing values from *N*(0, 1). Scale the *i*th column by eiv_sv_*i*. Solve for positive α and β such that α+β 4 and α / (α + β) = fraction_eiv. For each EIV, determine the number of affected genes by drawing from a Beta(α,β) distribution and multiplying by n_features and rounding to the nearest whole number. Randomly select this number of genes and set the loadings of all other genes to 0 for this EIV. Multiply the EIVs design matrix by the EIVs loading matrix to obtain the EIVs contribution to expression.
  c. Case/control status: Form the case-control design matrix as the one-hot encoding of the case-control status for each sample (n_samples x 2). Simulate a matrix of case-control effects (2 x n_features) by drawing values from a *N* (0,status_sd). Multiply the case-control design matrix the case-control effect matrix to obtain the case-control contribution to expression.
  d. Residual variation: Form the random noise matrix (n_samples x n_features)representing residual unexplained variability by drawing from *N* (0, resid_sd).
  e. Sum the output matrices from 4a-4d to form the final expression matrix.

#### Performing DEA on synthetic datasets

Five DEA approaches were applied to each synthetic dataset, differing by how the EIVs are addressed. SVA was implemented for the last three approach using the *sva* R package^95^:

1. **Oracle:** this method assumes that all parameters, including batch effects and EIVs, are known. DEA is performed while controlling for these known confounders, serving as an upper bound for the theoretically optimal performance that could be achieved when all parameters are known. DEA formula for **Oracle**: *Gene Hxpression* ~ *Status* + *Batch* + *EIV*_1_ + *…* + *EIV*_*n_EIV*_ (When n_EIV = 0, there are no EIV terms in the formula.)
2. **No SV Adjustment**: this method performs DEA on the normalized expression data, adjusting only for batch effect and omitting any EIV adjustment. It serves as a baseline to assess the impact of omitting EIVs. DEA formula for **No SVs**: *Gene Expression ~ Stu*.*tws + Batch*
3. **SV adjustments using a known number of SVs**: this method performs DEA assuming that the true number of EIVs n_eiv is known, but the EIVs themselves are unknown and must be estimated. Hence, SVA is used to estimate n_eiv SVs, which are then incorporated along with batch effects in the model. This approach assesses the impact of including the true number of SVs on DEA. DEA Formula for SVA **Known Number of SVs**:

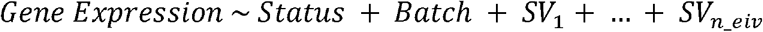

(When n_eiv = 0, there are no SV terms in the formula.)
4. **SV Adjustment with estimating the number of SVs using the “BE”** 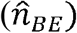 or (5) **“Leek”** 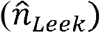 **methods**: these two methods perform DEA by using SVA to estimate a number of SVs determined by the BE and Leek methods respectively, and then incorporating all estimated SVs along with batch effects in the model. Compared to (3), these approaches assess the impact on DEA of estimating the number of SVs to be included in the model, which may result in using too many or too few SVs.

DEA Formula for **SVA “BE”**:

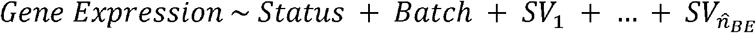

DEA Formula for **SVA “Leek”**:

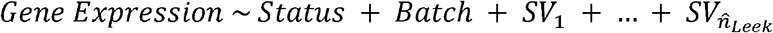

(When 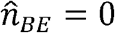 or 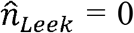, there are no SV terms in the formula, but note that as estimates, 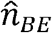 or 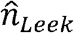 may be non-zero even when n_eiv = 0 and *vice versa*.)

DEA was performed using the lmFit function in the limma R package^96^. The features data matrices are transposed such that features are row and samples are columns. P-values were adjusted for multiple testing using the Benjamini-Hochberg method^97^, and genes with FDR ≤ 0.05 were considered DEGs. Performance metrics (TPR, FPR, FNR, TNR, FDR)^98, 99^ were calculated by comparing estimated DEGs from the DEA to the true DE status of each gene defined in the simulation metadata.

In total, there are 2,637,500 synthetic datasets were generated and analyzed using each of the five DEA approaches. Variance partition analyses^56^ confirmed that the synthetic data recapitulated variance components observed in real transcriptomic datasets.

### Evaluation of Disease Signal Correlation Maximization as a Strategy for FPR Control

A pair of gene expression datasets (Simulated Data 1 and Simulated Data 2) were simulated according to the above procedure (see “*Generating synthetic datasets*”) with one modification: the case-vs-control effect (step 4c) from Simulated Data 1 was reused in Simulated Data 2, such that both data sets shared the same disease signal but had independent noise structures. Each dataset had 500 subjects and 20,000 genes across 15 batches, incorporating batch effects, DEGs, and EIVs with specific parameters (n_subjects = 500, fraction_degs = 0.2, fraction_eiv = 0.4, batch_sd: 0.3, eiv_sd: 3, residual_sd: 0.2). For each dataset, DEA was performed multiple times adjusting for increasing numbers of estimated SVs (0 to 25, step of 1). For each SV number, this process was repeated 100 times with different random seeds, yielding 100 independent pairs of simulations per parameter condition. Across all repetitions, Spearman correlations of log_2_FC between datasets and true log_2_FC values were used to quantify cross-dataset reproducibility, while TPR and FPR assessed performance within each dataset.

### Real Brain Gene Expression Datasets

#### Data Description

Fourteen post-mortem human brain gene expression datasets were used to compare DEA results obtained with and without SVA across five neuropsychiatric disorders: AD, ASD, BP, SZ, and PD.

RNA-seq data were analyzed for AD, ASD, BP, and SZ, and microarray data were analyzed for PD. For AD, two datasets from the Accelerating Medicines Partnership - Alzheimer’s Disease (AMP-AD) consortium were used: the Religious Orders Study and Memory and Aging Project (ROSMAP)^62, 100^ and Mount Sinai/JJ Peters VA Medical Center Brain Bank (MSBB–Mount Sinai NIH Neurobiobank, herein MSBB)^101^. The ASD, BP, and SZ datasets were obtained from the PsychENCODE consortium. For ASD, two datasets were included: UCLA-ASD^48^ and Yale-ASD^44^. For BP and SZ, two datasets from the CommonMind Consortium (CMC) were used: the National Institute of Mental Health’s Human Brain Collection Core (NIMH-HBCC, herein CMC-HBCC)^18, 102^ and a dataset from the Mount Sinai School of Medicine, University of Pennsylvania, and University of Pittsburgh Brain Banks (herein CMC)^102^. For PD, eight datasets from the Gene Expression Omnibus (GEO) were included, with accession numbers GSE8397^67, 103^, GSE7621^68^, GSE24378^69^, GSE20292^69, 70^, GSE20141^69^, GSE20163^69^, GSE20164^69^, GSE49036^71^). These datasets were chosen based on a meta-analysis of PD postmortem brain transcriptomic study^104^. A subset of each dataset was analyzed in this study and details regarding number of samples are available in **Supplementary Table 2**.

#### Expression Data Processing

Quality control was performed on all brain RNA-seq and microarray datasets prior to DEA, both with and without SVA. For RNA-seq datasets, lowly expressed genes were removed following the filtering criteria defined in their respective studies (described below), normalized using Trimmed Mean of M-values via calcNormFactors function from the edgeR R package^105^, and transformed to log2-CPM with precision weights using voom function from the limma R package^96^ prior to linear modeling. Microarray datasets underwent log_2_ transformation and quantile normalization, following previous protocol^106^. The resulting number of genes and sample sizes for each dataset is in **Supplementary Table 2**. The following normalization formulas were used, matching those applied in studies performing DEA on these datasets. Regression formulas were selected based on prior literature, and for RNA-seq datasets, the same formula was used for both voom and DEA.

##### Dataset 1: AD - ROSMAP^5^

The formula is:

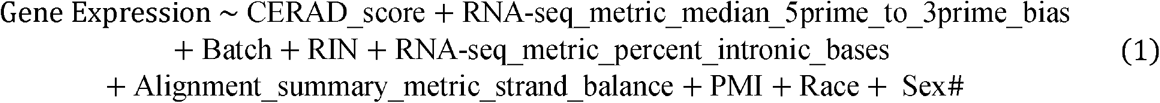

where CERAD_score is the Consortium to Establish a Registry for Alzheimer’s Disease (CERAD) score^107^, which was used to define case–control status (CERAD score = 1, “ceradsc1,” as AD; CERAD score = 4, “ceradsc4,” as control); RNA-seq_metric_median_5prime_to_3prime_bias is the median 5’ to 3’ bias; Batch is batch effects; RIN is RNA Integrity Number; RNA-seq_metric_percent_intronic_bases is the percentage of intronic bases; Alignment_summary_metric_strand_balance is strand balance; PMI is post-mortem interval; Race is reported race; Sex is reported sex. While some of these variables were modeled as random effects in previous works, they are treated as fixed effect using limma in this study. Genes with ≤ 1 count per million (CPM) in at least 10% of samples were excluded.

##### Dataset 2: AD - MSBB^5^

The formula is:

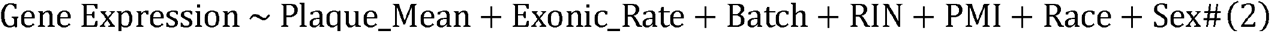

where Plaque_Mean is the mean neocortical amyloid-β plaque density (number of plaques/mm^2^);

Exonic_Rate is the fraction of reads that map within exons; Batch is batch effects; RIN is RNA Integrity Number; Race is reported race; Sex is reported sex. While some of these variables were modeled as random effects in previous works, they are treated as fixed effect using limma in this study. Genes with ≤ 1 CPM in at least 10% of samples were excluded.

##### Dataset 3: BP&SZ – CMC-HBCC^18^

The formula is:

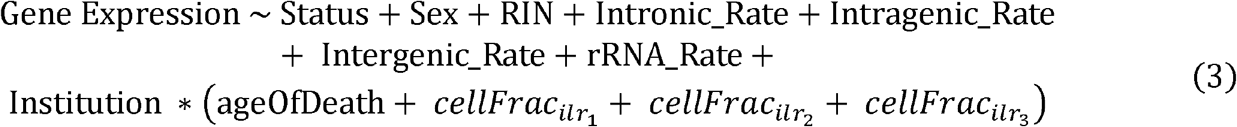

where Status is a diagnosis of SZ or control; Sex is reported sex; Intronic_Rate is the fraction of reads that map within introns; Intragenic_Rate is the fraction of reads that map within genes (within introns or exons); Intergenic_Rate is the fraction of reads that map to the genomic space between genes; rRNA_Rate is the ratio of all reads aligned to rRNA regions to total reads; Institution is the research site where data was collected (MSSM, Penn, or Pitt); ageOfDeath is the participant’s age at death; 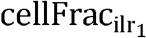, 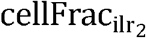, 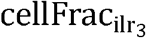 are the transformed isometric log ratio of cell type composition. Genes with ≤ 0.5 CPM in at least 30% of samples were excluded.

##### Dataset 4: BP&SZ - CMC^18^

The same processing approach and voom normalization formula described for HBCC were applied. The only difference here is that Diagnosis refers to a diagnosis of BP or control. Genes with ≤ 0.5 CPM in at least 30% of samples were excluded.

##### Dataset 5: ASD – UCLA-ASD^108^

The formula is:

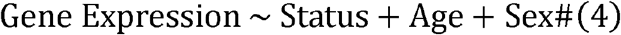

where Status is a diagnosis of ASD or control; Age is self-reported age; Sex is self-reported sex. Genes with ≤ 1 CPM in at least 10% of samples were excluded.

##### Dataset 6: ASD – Yale-ASD

The same formula described for UCLA-ASD were applied^108^. Genes with ≤ 1 CPM in at least 10% of samples were excluded.

##### Datasets 7-14: PD – GEOs

The formula for **GSE8397**^67, 103^ is:

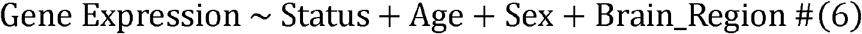

where Status is a diagnosis of PD or control; Age is self-reported age; Sex is self-reported sex; Brain_Region refers to brain regions (medial substantia nigra or superior frontal gyrus). This dataset was generated using the GPL96 microarray platform.

The formula for **GSE7621**^68^ is:

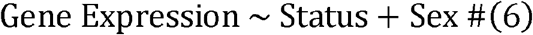

where Status is a diagnosis of PD or control; Sex is self-reported sex. This dataset was generated using the GPL570 microarray platform.

The formula for **GSE24378**^69^ is:

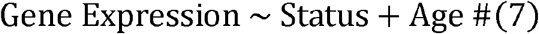

where Status is a diagnosis of PD or control; Age is self-reported age. This dataset was generated using the GPL1352 microarray platform.

The formula for **GSE20292**^69, 70^ is:

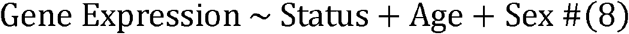

where Status is a diagnosis of PD or control; Age is self-reported age; Sex is self-reported sex.

This dataset was generated using the GPL96 microarray platform.

The formula for **GSE20141**^69^ is:

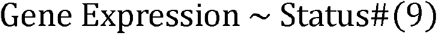

where Status is a diagnosis of PD or control. This dataset was generated using the GPL570 microarray platform.

The formula for **GSE20163**^69^ is:

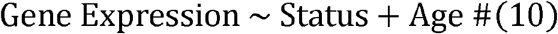

where Status is a diagnosis of PD or control; Age is self-reported age. This dataset was generated using the GPL96 microarray platform.

The formula for **GSE20164**^69^ is:

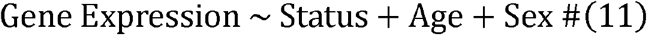

where Status is a diagnosis of PD or control; Age is self-reported age; Sex is self-reported sex.

This dataset was generated using the GPL96 microarray platform.

The formula for **GSE49036**^71^ is:

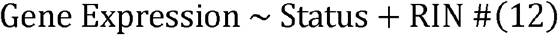

where Status is a diagnosis of PD or control; RIN is RNA integrity number. This dataset was generated using the GPL570 microarray platform.

#### DEA Methods for Real Data

After gene expression data processing, DEA was performed using the same approach as in the simulation. Specifically, DEA was performed using lmFit function in limma^96^ on the normalized gene expression data adjusted for covariates listed in the aforementioned formula for specific datasets. P-values were adjusted for multiple testing using the Benjamini-Hochberg method^97^, and genes with FDR ≤ 0.05 were considered differentially expressed.

#### SVA for Real Data

SVA was run on each dataset using the sva R package^95^, employing both “BE” and “Leek” methods to estimate the number of SVs. A full model matrix (including the primary variable of interest and other known covariates) and a null model matrix (including only the known covariates) were constructed. The null model in SVA enables the identification of variation that is independent of the primary variable of interest^36^. The identified SVs were then included as additional covariates in the limma-voom DEA, alongside the known covariates (as specified in the voom regression formulas). The number of SVs identified by each method varied across datasets (**Supplementary Table 2**).

#### Assessment of DE signatures reproducibility with and without SVA

Reproducibility of DE signatures was quantified by calculating Spearman rank correlations of log_2_FC values between dataset pairs for AD, ASD, BP, and SZ, and between all distinct pairs among the eight PD datasets. To assess the impact of incorporating a small number of SVs on the reproducibility of DE signatures, Spearman correlation of log_2_FC was calculated between the ROSMAP and MSBB datasets for each number of SVs, ranging from 0 to the maximum number of SVs identified by the SVA “BE” method (**Supplementary Table 2**).

External reproducibility of DE signatures was assessed by testing their overlap with published DEG lists. We performed one-sided Fisher’s exact tests for enrichment between our DEGs (with and without SVA) and published DEG lists from 9 independent external datasets in AD^74–82^, 5 in BP^10, 83–86^, and 5 in SZ^87–90^. We then used one-sided paired Wilcoxon signed-rank tests (alternative = “less”) against our DEG results and the resulting p-values to compare the significance of DEG overlap between the SVA and non-SVA approaches for each dataset.

#### Cosine Similarity Analysis

To evaluate the angles between SVs and the primary variable of interest, we calculated the cosine similarity between columns of the design matrix and the EIV matrix (the dot product of each column pair divided by the product of their Euclidean norms), resulting in a cosine similarity matrix. These values (clamped between −1 and 1) were converted to angles (degrees) using the inverse cosine function, taking the supplement of angles >90° to represent the minimum angular separation.

#### Evaluating the Impact of SVA on DEA under Null Conditions

To assess SVA impact on DEG distributions under null conditions where the true biological signals are removed, we shuffled group labels (e.g., “ceradsc1”, “ceradsc4”) in ROSMAP and performed DEA with and without SVA. The SVA pipeline incorporated 0 to 49 SVs, while the non-SVA pipeline considered only shuffled labels and batch effects. Using limma, we recorded the number of DEGs (FDR < 0.05) across 100 shuffles for each group.

#### Other statistical analyses

Data processing and statistical analysis was performed using the R 4.2.0.^109^ and R packages from Bioconductor^110, 111^. Parallel and distributed computing were performed using BiocParallel (v1.32.6)^111^ and foreach (v 1.5.2)^112^. Gene annotation and ID conversion were performed using biomaRt (v2.54.1)^113^. Publicly available gene expression datasets were retrieved and processed using GEOquery (v2.70.0)^114^. Large tables were read-in and processed using many tidyverse^115^ packages and using the R package data.table (v1.16.0)^116^ and readr (v2.1.5)^117^. Data integrity and logical validation were ensured using assertthat (v0.2.1). Data wrangling, iteration, and reshaping were conducted using dplyr (v1.0.10), purrr (v1.0.4), tibble (v3.2.1), and tidyr (v1.3.1). Figures were generated using the R package ggplot2 (v3.5.1)^118^, ggrastr (v1.0.2)^119^, ggrepel (v0.9.6)^120^, viridis (v0.6.5)^121^, and patchwork (v1.3.0)^122^. For heatmap visualization and cluster ordering, pairwise distances were computed via dist function in stats (v.4.2.0)^123, 124^, and rows and columns were reordered via seriation (v.1.5.6)^125^. Differential expression analysis was conducted using limma (v3.54.2)^96^, edgeR (v3.40.2)^105^, and sva (v3.46.0)^95^, while variance decomposition was analyzed using variancePartition^56^. Spearman correlations, Fisher’s exact tests, and one-sided paired Wilcoxon signed-rank tests were run with cor.test() (alternative = “greater”), fisher.test() (alternative = “greater”), and wilcox.test() (alternative = “less”) functions, respectively, in the stats package^109^. Multiple testing correction was performed using Benjamini-Hochberg procedure using the p.adjust R function^97^.

## Supporting information

Supplementary Table 1_SVABE

Supplementary Table 1_Oracle

Supplementary Table 1_NoSVA

Supplementary Table 1_KnownNumberofSVs

Supplementary Table 1_SVALeek

Supplementary Table 2_ASD

## Extended Data Figure legends

**Extended Data Fig. 1:**
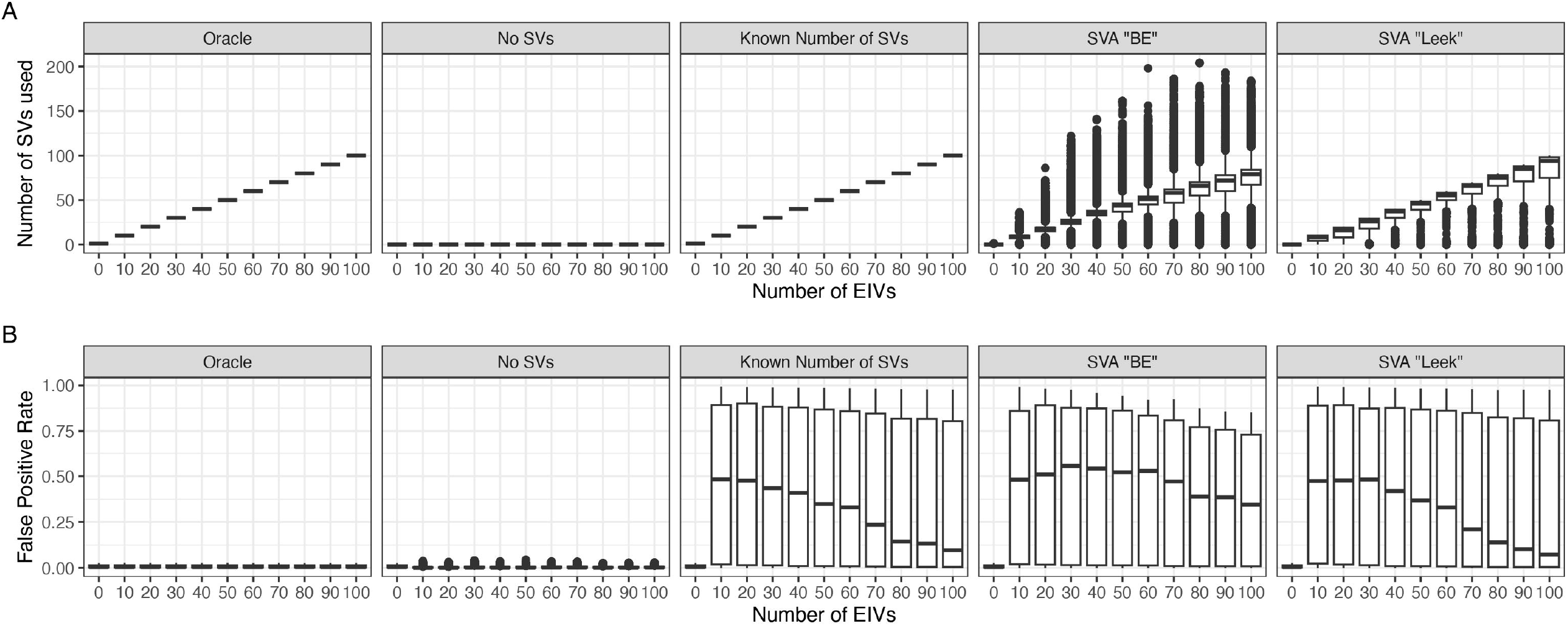
Methods using SVs generally estimate the numbers of EIVs accurately. (**A**) Boxplots comparing the number of EIVs (x-axis) to the number of SVs used by each approach (y-axis). (**B**) Boxplot showing the relationship between the number of EIVs (x-axis) and FPR (y-axis) for each approach.

**Extended Data Fig. 2:**
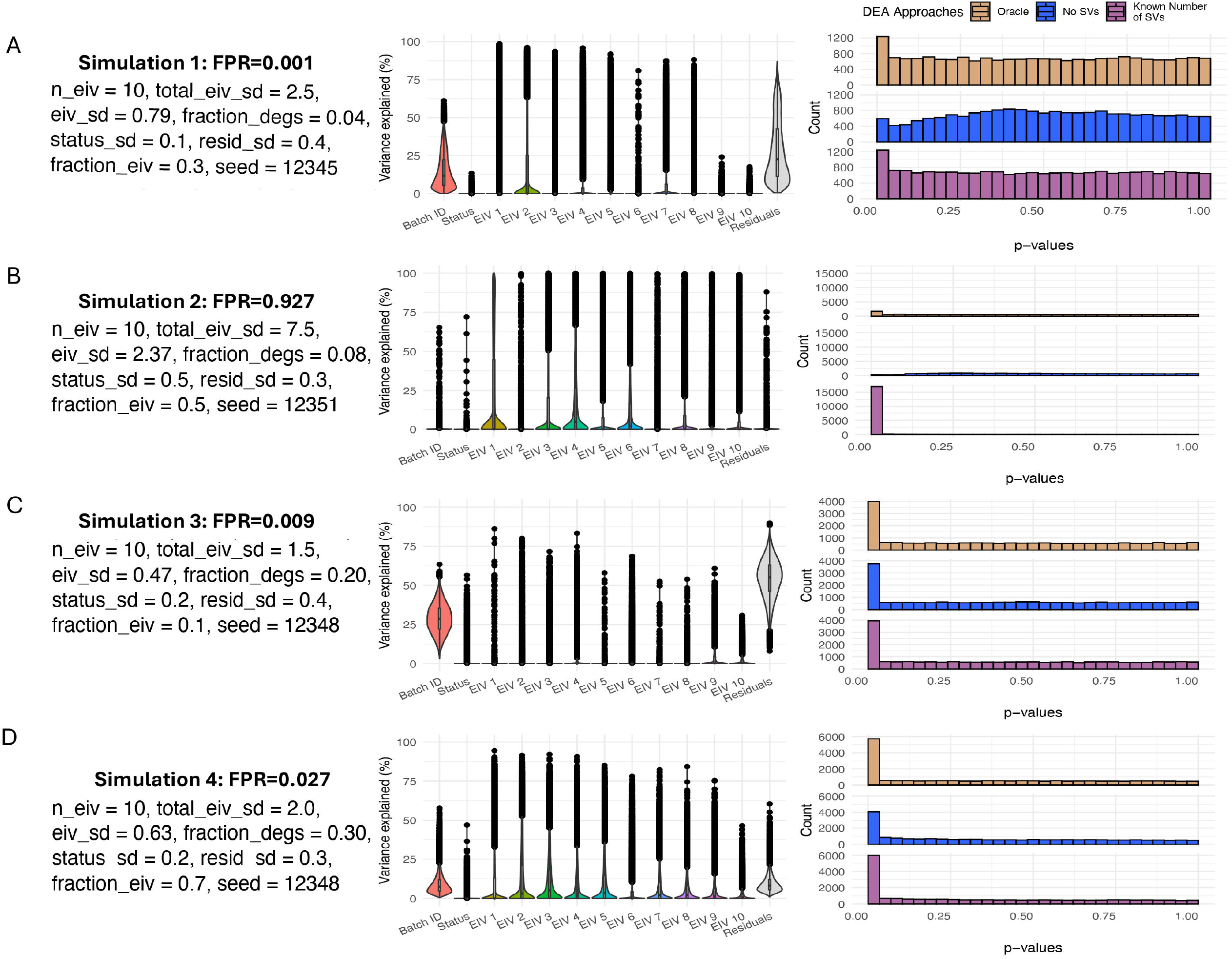
Variance partitioning and p-value distributions for four representative simulations. Variance decomposition and p-value distributions for four randomly selected representative simulations with EIVs = 10 are displayed. Each panel (**A-D**) displays: simulation parameters and the resulting false positive rate FPR; variance partitioning showing the percentage of variance explained (y-axis) by disease status and each covariate across all genes (x-axis); p-value distributions comparing three DEA approaches (defined in the legend). The x-axis is the p-values; the y-axis is the count of each p-value. The simulation parameters are: n_eiv = number of EIVs; total_eiv_sd = total EIV variation; eiv_sd = standard deviation per EIV; fraction_degs = fraction of differentially expressed genes; status_sd = disease status effect size; resid_sd = residual standard deviation; fraction_eiv = fraction of genes affected per EIV; seed = random seed for reproducibility.

**Extended Data Fig. 3:**
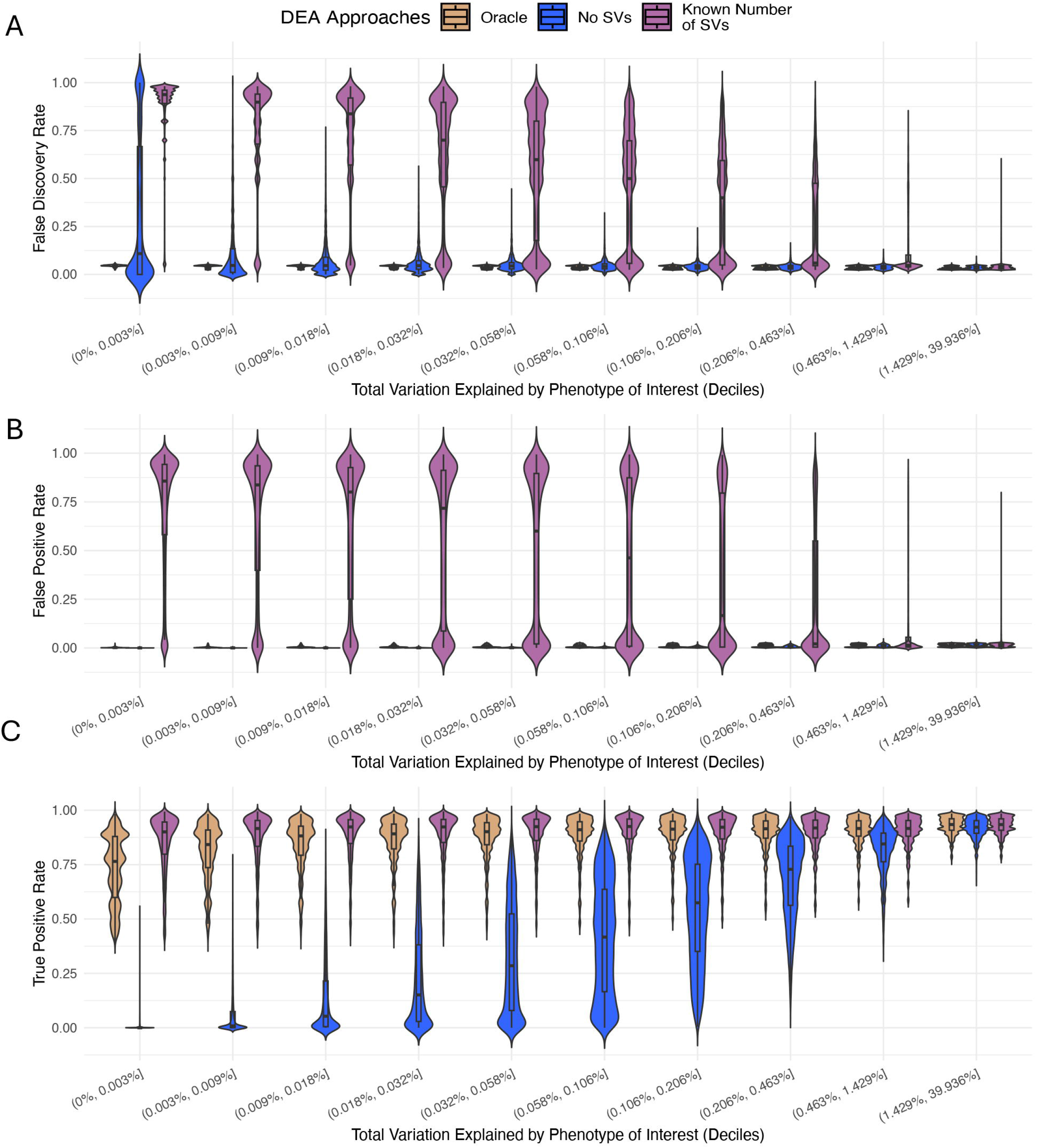
Stronger signal strengths correspond to lower FDR, FPR, and do not affect TPR for SVA Known Number of SVs. Violin and boxplots displaying the results from all simulations across three DEA approaches (defined in the legend). Synthetic data are stratified by signal strength, defined as the total variance explained by the phenotype of interest (i.e., disease status) and grouped into deciles. The x-axis is the percent variance explained by the phenotype of interest within each decile; the y-axis is (**A**) FDR, (**B**) FPR, and (**C**) TPR, respectively. For TPR, “Oracle” and “SVA Known Number of SVs” approaches consistently achieved near perfect TPR across phenotype signal strength (**C**), while the TPR of the “No SVs” approach improved markedly as the phenotype signal strength increased (**C**). On the other hand, these methods showed contrasting patterns for false discovery control. “Oracle” and “No SVs” approaches maintained low FPR (**B**) and FDR (**A**), whereas “SVA Known Number of SVs” frequently exhibited uncontrolled FPR (**B**) and FDR (**A**), a concerning pattern that diminished only as phenotype signal strength increased.

**Extended Data Fig. 4:**
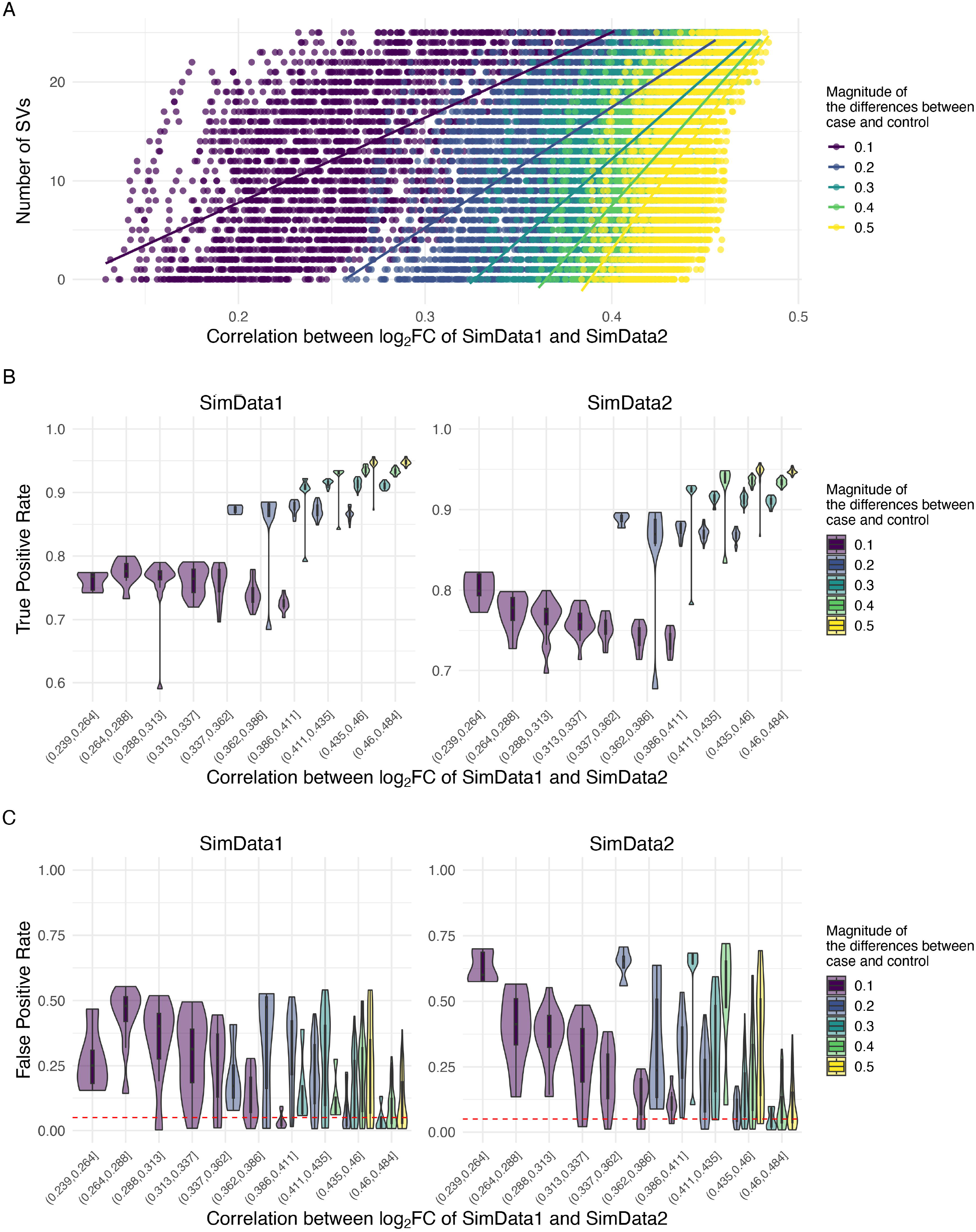
Relationships of the correlation of log_2_FC between independent simulated datasets and number of SVs, TPR, and FPR by signal strength. Two simulated datasets (SimData1 and SimData2) were generated with identical signal and number of true DEGs but different noise structures. (**A**) Scatterplot showing the correlation of log_2_FC between SimData1 and SimData2 (x-axis) against the numbers of SVs (y-axis). Points are colored by the magnitude of the case–control effect size, and the fitted line (linear model) depicts the relationship between correlation and number of SVs. (**B**-**C**) Violin and boxplots showing the correlation of log_2_FC between SimData1 and SimData2 (x-axis) against their respective TPR (**B**) and FPR (**C**) (y-axis), faceted by each simulated dataset.

**Extended Data Fig. 5:**
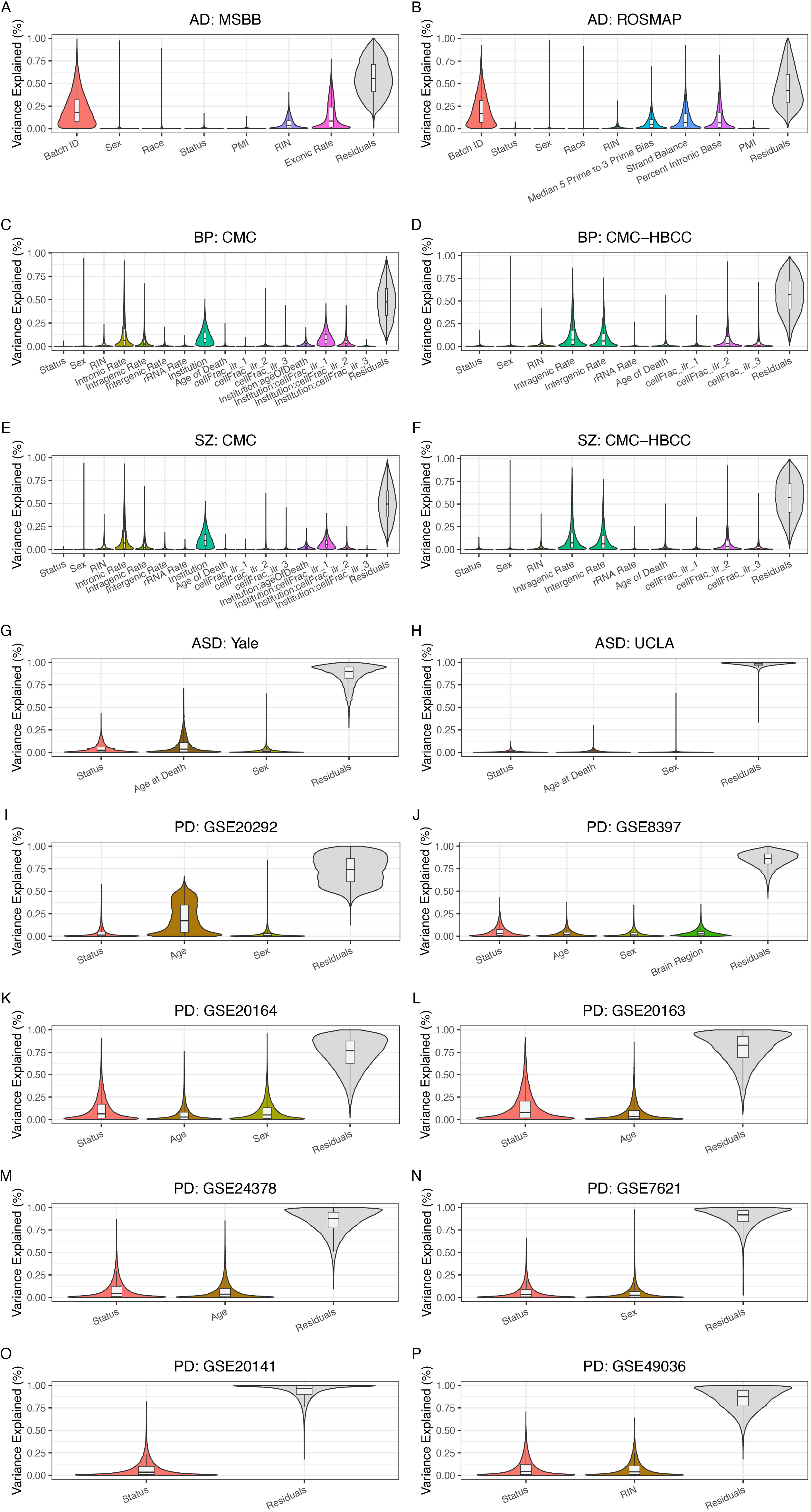
Variance partition plots of six real neuropsychiatric gene expression datasets. Violin and boxplots showing the distribution of percent of variance explained (y-axis) by the phenotype of interest (i.e., status for disease status) and covariates across genes (y-axis) for six neuropsychiatric datasets: **AD**: (**A**) ROSMAP and (**B**) MSBB; **BP**: (**C**) CMC and (**D**) CMC-HBCC; SZ: (**E**) CMC and (**F**) CMC-HBCC.

**Extended Data Fig. 6:**
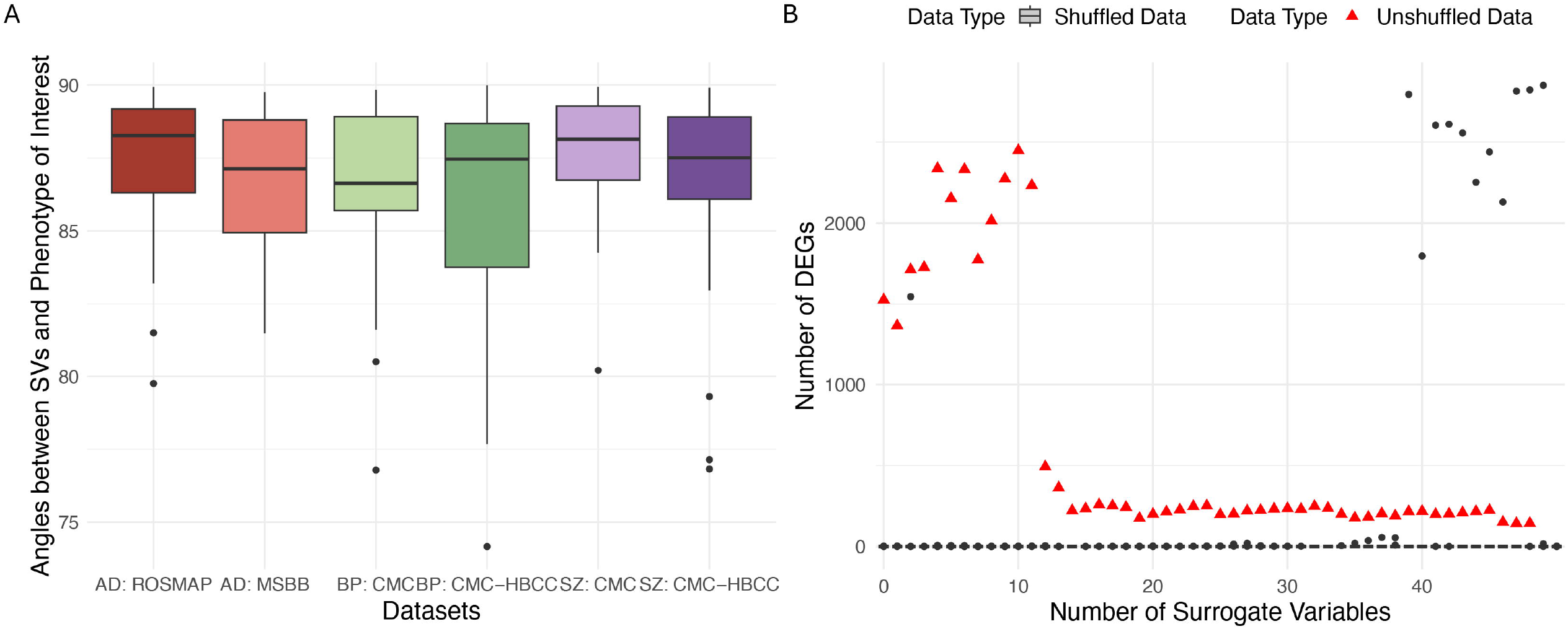
Cosine angle between SVs and the phenotype of interest, and relationship between number of SVs and number of DEGs when phenotype is not present in ROSMAP. (**A**) Boxplots showing the distribution of the cosine angles (x-axis) between estimated SVs and phenotype (i.e. disease status) across six neuropsychiatric datasets: **AD**: ROSMAP and MSBB; **BP and SZ**: CMC and CMC-HBCC (y-axis). (**B**) Boxplots showing the number of DEGs detected (y-axis) when using increasing numbers of SVs (x-axis) under null conditions where the phenotype is not present via shuffling the case labels. The null condition is in grey boxes; the true unshuffled case-control status is in red triangles.

**Extended Data Fig. 7:**
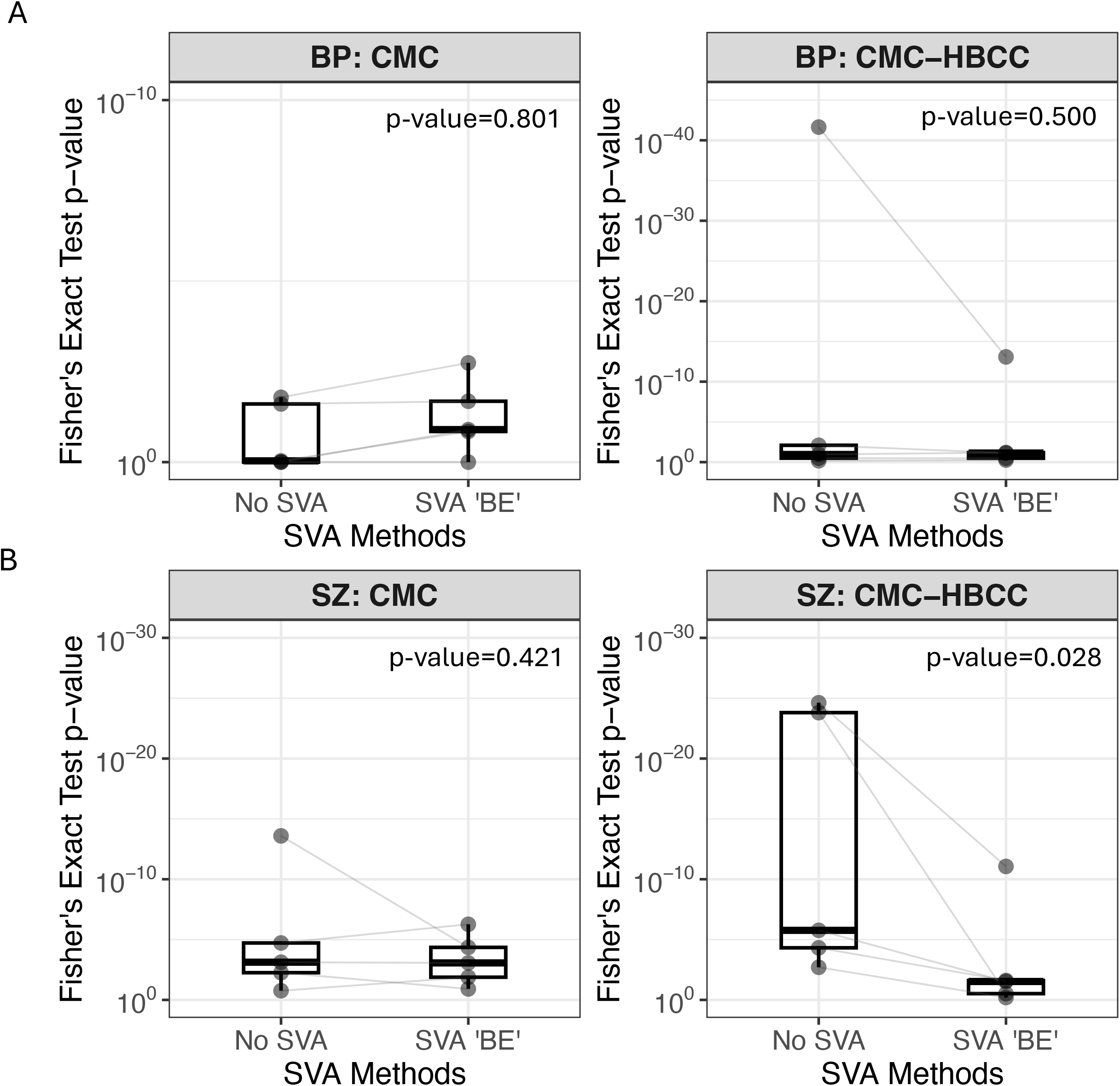
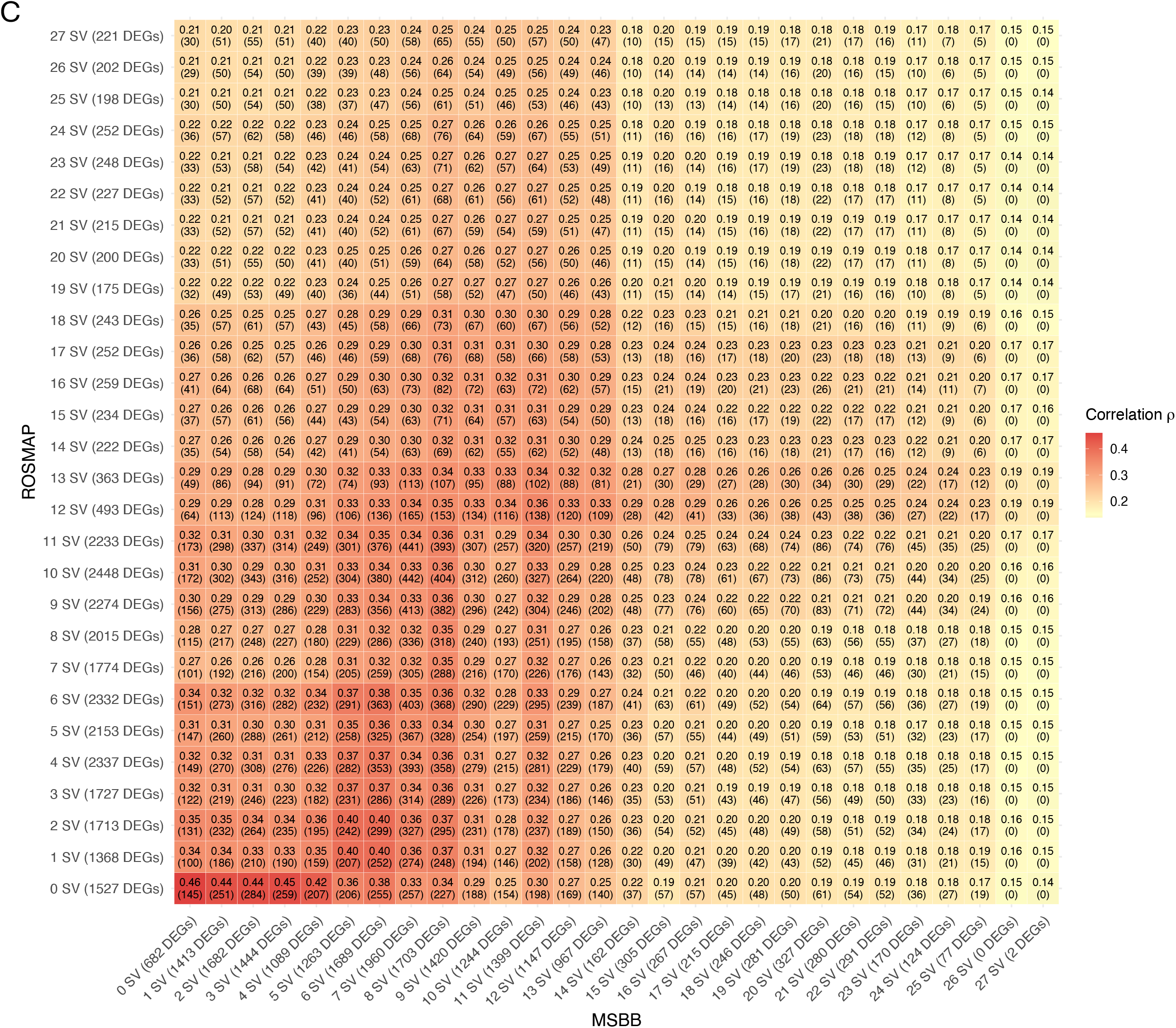
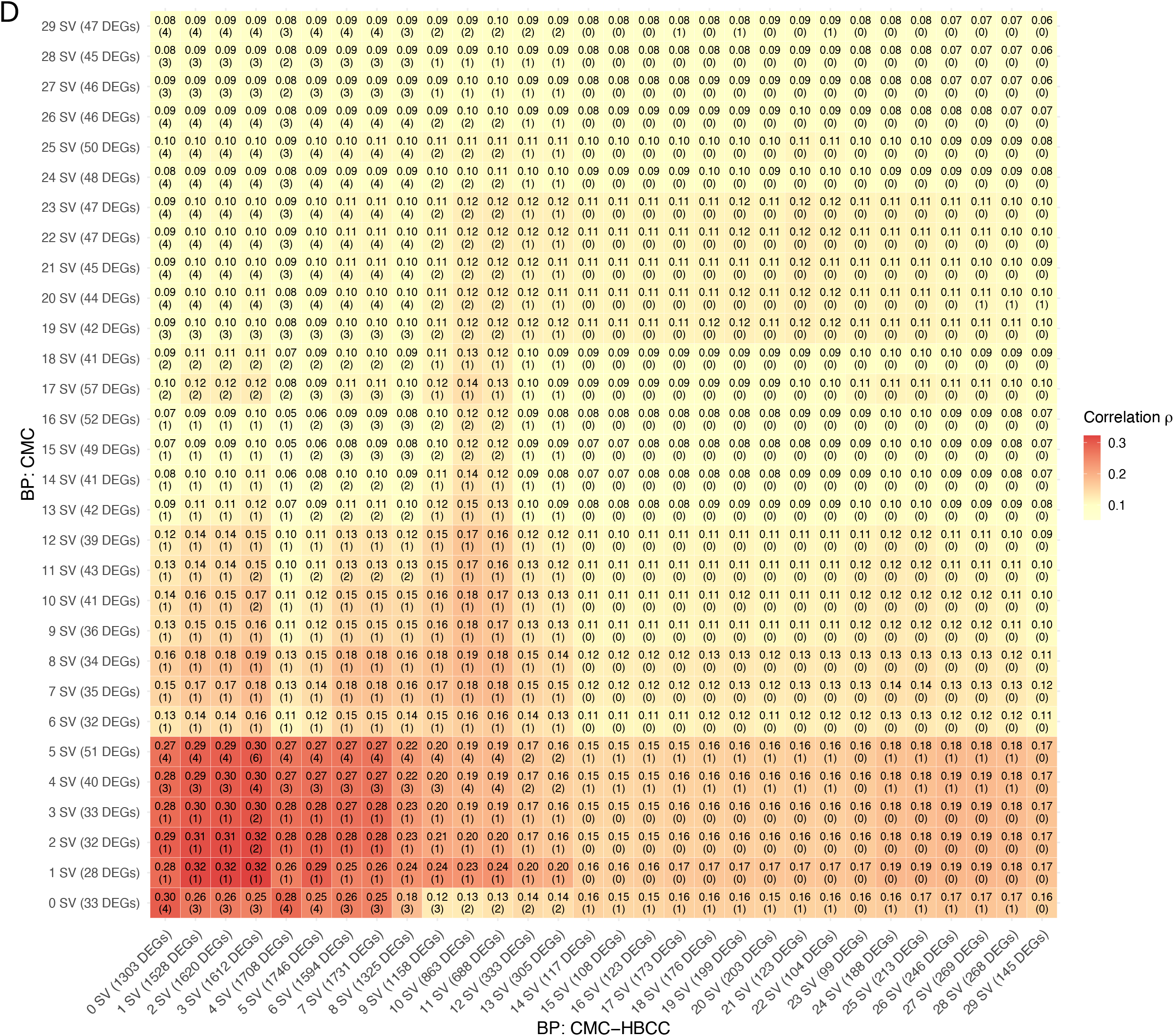
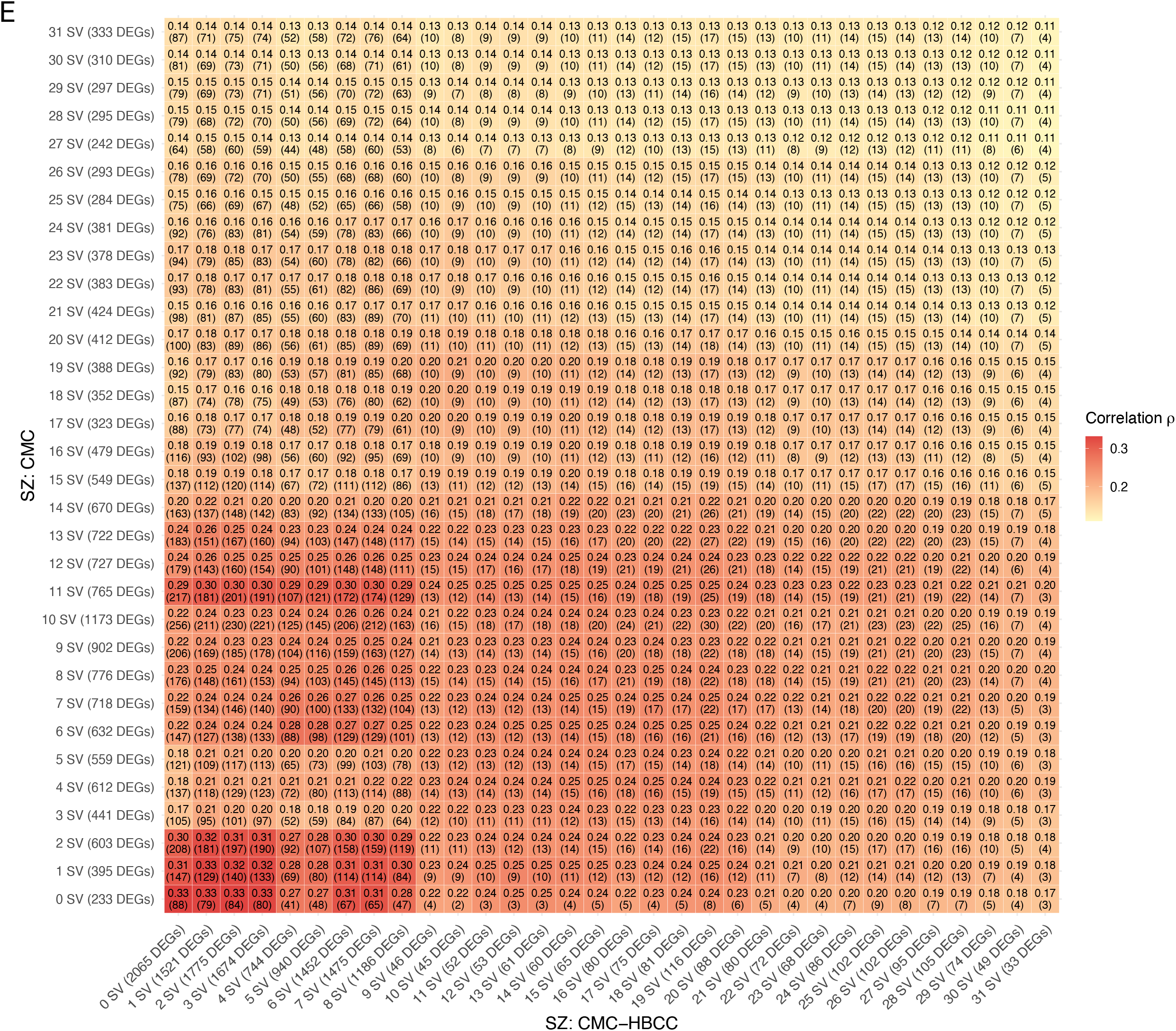
SVA Reduces Reproducibility of BP and SZ DE Signatures. (**A-B**) Box plots comparing the statistical significance of DEG overlap between two primary cohorts (CMC and CMC-HBCC) and five additional publicly available datasets for BP and SZ, respectively. The x-axis shows two methods: No SVA (left) and SVA “BE” (right), where SVA “BE” estimated 29 SVs for BP: CMC and 36 SVs for BP: CMC-HBCC (**A**), and 50 SVs for SZ: CMC and 13 SVs for SZ: CMC-HBCC (**B**); the y-axis shows negative log-transformed p-values from Fisher’s exact tests, where higher values indicate more significant overlap (better reproducibility). The FDR-corrected p-values from Wilcoxon signed-rank tests are displayed in each facet. (**C-E**) Heatmap showing Spearman correlations of log_2_FC between dataset pairs: ROSMAP vs MSBB (**C**), BP: CMC vs BP: CMC-HBCC (**D**), and SZ: CMC vs SZ: CMC-HBCC (**E**) when incrementally including SVs (0-10 SVs). Each cell displays the Spearman ρ values with the number of overlapping significant DEGs in parentheses. Higher Spearman ρ and more overlapping DEGs indicate better reproducibility between datasets. SVA estimated maximum SVs of 27 for MSBB, 29 for BP: CMC-HBCC, and 31 for SZ: CMC-HBCC.

**Extended Data Fig. 8:**
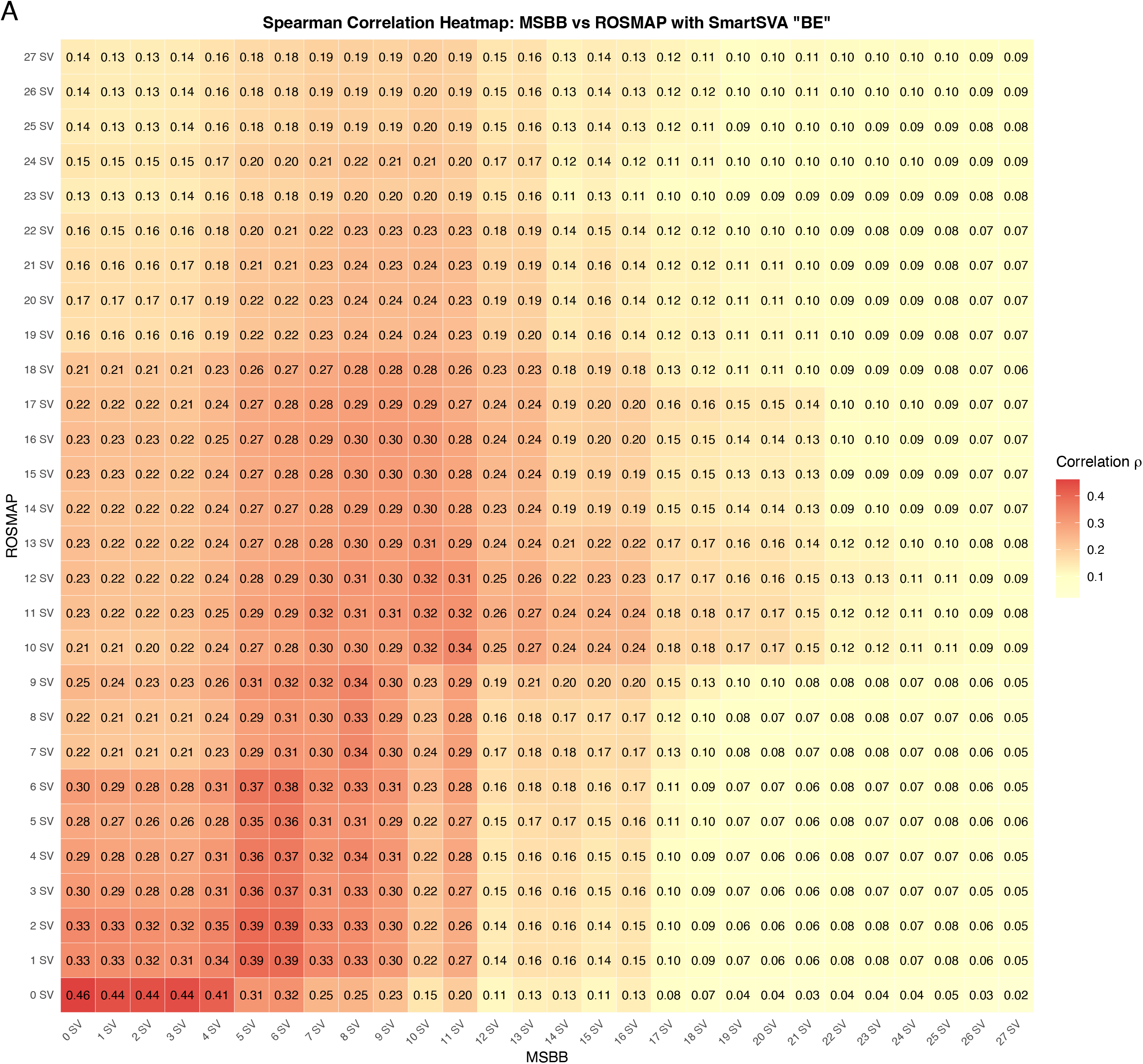
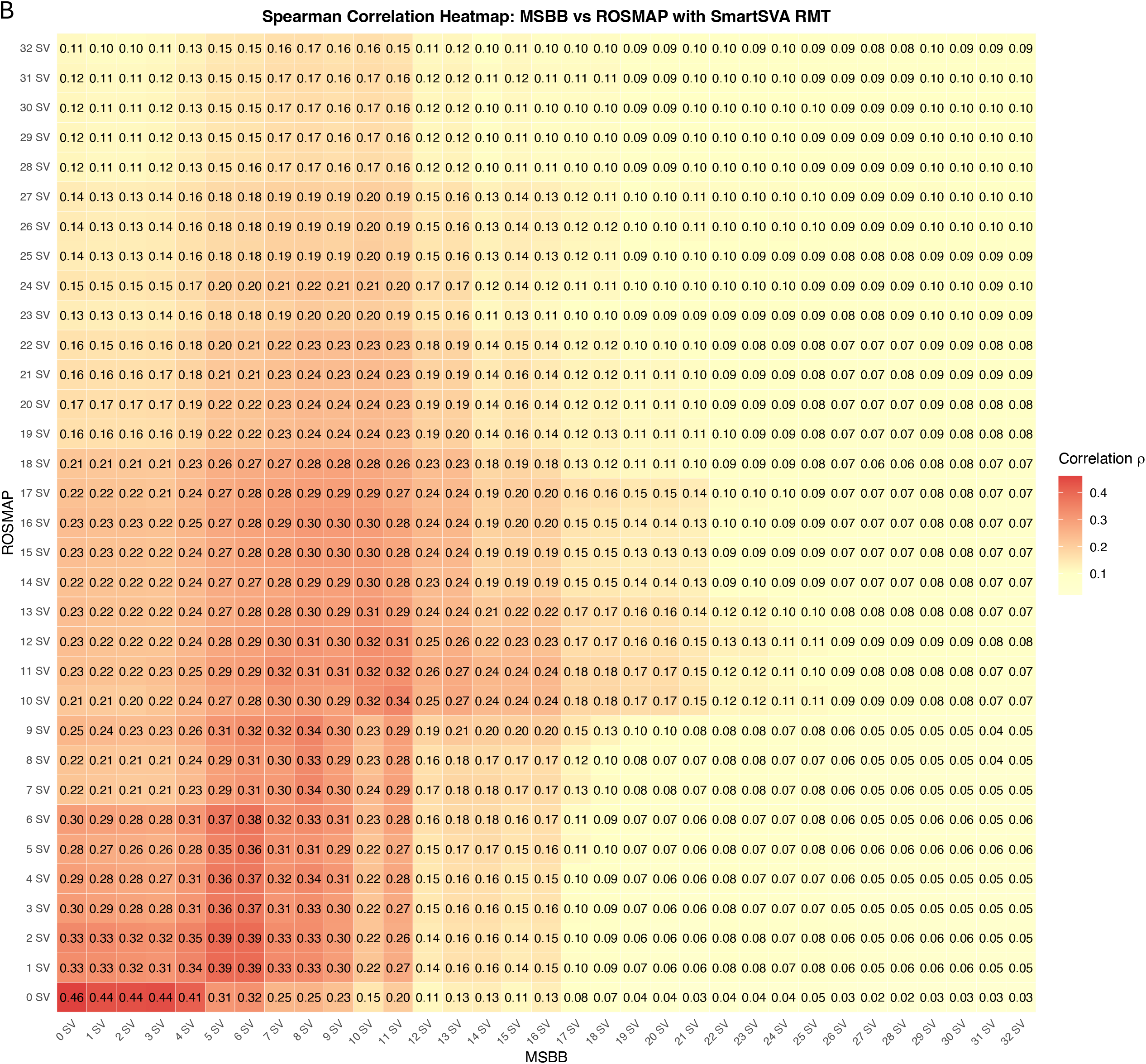
Convergence-based SVA Also Reduces Reproducibility of AD DE Signatures. (**A-B**) Heatmap showing Spearman correlations of log_2_FC between ROSMAP (x-axis) and MSBB (y-axis) when incrementally including SVs (0-10 SVs) using SmartSVA “BE” (**A**) and SmartSVA “Random Matrix Theory (RMT)” (**B**). Each cell displays the Spearman ρ values. Higher Spearman ρ and more overlapping DEGs indicate better reproducibility between datasets. The “BE” method selected 27 SVs for MSBB and 47 SVs for ROSMAP, while RMT selected 32 SVs for MSBB and 68 SVs for ROSMAP.

## Description of Supplemental Tables

**Supplementary Table 1:** Simulation parameters along with their true positive rate, true negative rate, false positive rate, false negative rate, and mean, median, and standard deviation of variation of status.

**Supplementary Table 2:** Real datasets characteristics and all their differential expression analyses results with and without SVA adjustments.

## Acknowledgements

We thank the departments of Artificial Intelligence of Human Health, Neuroscience, Medicine, and Psychiatry at the Icahn School of Medicine at Mount Sinai for their support in making this work possible. This work was supported in part through the computational and data resources and staff expertise provided by Scientific Computing and Data at the Icahn School of Medicine at Mount Sinai and supported by the Clinical and Translational Science Awards (CTSA) grant UL1TR004419 from the National Center for Advancing Translational Sciences. Special thanks to Charles Deng for providing critical feedback and helping with the visualization of schematics for the manuscript.

## Data Availability Statement

All data used are publicly available. The AD datasets (MSBB and ROSMAP) are available at https://adknowledgeportal.synapse.org/Explore/Studies/DetailsPage?Study=syn3159438 and https://adknowledgeportal.synapse.org/Explore/Studies/DetailsPage?Study=syn3219045, respectively. The ASD, BP, and SZ datasets are available through PscyhENCODE Capstone Collection and can be accessed at http://www.doi.org/10.7303/syn12080241. The PD datasets are publicly available through GEO at https://www.ncbi.nlm.nih.gov/geo/; GSE8397:

https://www.ncbi.nlm.nih.gov/geo/query/acc.cgi?acc=GSE8397; GSE7621:

https://www.ncbi.nlm.nih.gov/geo/query/acc.cgi?acc=GSE7621; GSE24378:

https://www.ncbi.nlm.nih.gov/geo/query/acc.cgi?acc=GSE24378; GSE20292:

https://www.ncbi.nlm.nih.gov/geo/query/acc.cgi?acc=GSE20292; GSE20141:

https://www.ncbi.nlm.nih.gov/geo/query/acc.cgi?acc=GSE20141; GSE20163:

https://www.ncbi.nlm.nih.gov/geo/query/acc.cgi?acc=GSE20163; GSE20164:

https://www.ncbi.nlm.nih.gov/geo/query/acc.cgi?acc=GSE20164; GSE49036:

https://www.ncbi.nlm.nih.gov/geo/query/acc.cgi?acc=GSE49036.

## Code Availability

All code required to reproduce the analyses and generate figures presented in this manuscript is publicly available at https://github.com/BeckmannLab/sva_eval_dea_neuropsych.

## Author Contributions

The following authors contributed to the overall conception of research idea and study design: N.D.B.. The following authors performed all analyses and wrote the manuscript: N.D.B. and D.T.H.. The following authors contributed to data processing and analyses: N.D.B., D.T.H., R.C.T., and A.N.L. The following authors contributed to preparing figures and tables: N.D.B., D.T.H., A.N.L., R.C.T., N.G., E.K., D.J., and A.S.B.. The following authors provided critical feedback to writing and analyses results: R.C.T., A.N.L., N.G., A.W.C., G.N.N..

## References

1. Begley, C.G. & Ioannidis, J.P. Reproducibility in science: improving the standard for basic and preclinical research. Circulation research 116, 116–126 (2015).

2. Kapoor, S. & Narayanan, A. Leakage and the reproducibility crisis in machine-learningbased science. Patterns 4 (2023).

3. Baker, M. (Nature Publishing Group UK London, 2016).

4. Baykal, P.I. et al. Genomic reproducibility in the bioinformatics era. Genome Biology 25, 213 (2024).

5. Beckmann, N.D. et al. Multiscale causal networks identify VGF as a key regulator of Alzheimer’s disease. Nature Communications 11 (2020).

6. Raj, T. et al. Integrative transcriptome analyses of the aging brain implicate altered splicing in Alzheimer’s disease susceptibility. Nat Genet 50, 1584–1592 (2018).

7. Annese, A. et al. Whole transcriptome profiling of Late-Onset Alzheimer’s Disease patients provides insights into the molecular changes involved in the disease. Sci Rep 8, 4282 (2018).

8. Stott, J. et al. A systematic review of non-coding RNA genes with differential expression profiles associated with autism spectrum disorders. PLOS ONE 18, e0287131 (2023).

9. Gandal, M.J. et al. Broad transcriptomic dysregulation occurs across the cerebral cortex in ASD. Nature 611, 532–539 (2022).

10. Park, S.W., Seo, M.K., Webster, M.J., Lee, J.G. & Kim, S. Differential expression of gene co-expression networks related to the mTOR signaling pathway in bipolar disorder. Transl Psychiatry 12, 184 (2022).

11. Nurnberger, J.I. et al. Identification of pathways for bipolar disorder: a meta-analysis. JAMA psychiatry 71, 657–664 (2014).

12. Pacifico, R. & Davis, R. Transcriptome sequencing implicates dorsal striatum-specific gene network, immune response and energy metabolism pathways in bipolar disorder. Molecular psychiatry 22, 441–449 (2017).

13. Quan, W. et al. Identification of potential core genes in Parkinson’s disease using bioinformatics analysis. Parkinson’s Disease 2021, 1690341 (2021).

14. Borrageiro, G., Haylett, W., Seedat, S., Kuivaniemi, H. & Bardien, S. A review of genome-wide transcriptomics studies in Parkinson’s disease. European Journal of Neuroscience 47, 1–16 (2018).

15. Glaab, E. & Schneider, R. Comparative pathway and network analysis of brain transcriptome changes during adult aging and in Parkinson’s disease. Neurobiology of disease 74, 1–13 (2015).

16. Hauberg, M.E. et al. Differential activity of transcribed enhancers in the prefrontal cortex of 537 cases with schizophrenia and controls. Molecular psychiatry 24, 1685–1695 (2019).

17. Merikangas, A.K. et al. What genes are differentially expressed in individuals with schizophrenia? A systematic review. Molecular psychiatry 27, 1373–1383 (2022).

18. Hoffman, G.E. et al. Sex differences in the human brain transcriptome of cases with schizophrenia. Biological psychiatry 91, 92–101 (2022).

19. Rosati, D. et al. Differential gene expression analysis pipelines and bioinformatic tools for the identification of specific biomarkers: a review. Comput Struct Biotechnol J (2024).

20. Xie, Z. et al. Identification of therapeutic targets for Alzheimer’s Disease Treatment using bioinformatics and machine learning. Sci Rep 15, 3888 (2025).

21. Lamisa, A.B. et al. A meta-analysis of bulk RNA-seq datasets identifies potential biomarkers and repurposable therapeutics against Alzheimer’s disease. Sci Rep 14, 24717 (2024).

22. Nieuwenhuis, T.O. et al. Patterns of unwanted biological and technical expression variation among 49 human tissues. Laboratory Investigation 104, 102069 (2024).

23. Yu, Y., Mai, Y., Zheng, Y. & Shi, L. Assessing and mitigating batch effects in largescale omics studies. Genome Biology 25, 254 (2024).

24. Hamaguchi, Y., Zeng, C. & Hamada, M. Impact of human gene annotations on RNA-seq differential expression analysis. BMC Genomics 22, 1–12 (2021).

25. Leek, J.T. et al. Tackling the widespread and critical impact of batch effects in high-throughput data. Nat Rev Genet 11, 733–739 (2010).

26. Scherer, A. Batch effects and noise in microarray experiments: sources and solutions, Vol. 868. (Wiley Online Library, 2009).

27. McCarthy, D.J., Chen, Y. & Smyth, G.K. Differential expression analysis of multifactor RNA-Seq experiments with respect to biological variation. Nucleic acids research 40, 4288–4297 (2012).

28. Law, C.W., Chen, Y., Shi, W. & Smyth, G.K. voom: Precision weights unlock linear model analysis tools for RNA-seq read counts. Genome biology 15, 1–17 (2014).

29. Hui, H.W.H., Kong, W. & Goh, W.W.B. Thinking points for effective batch correction on biomedical data. Briefings in Bioinformatics 25, bbae515 (2024).

30. Stegle, O., Parts, L., Durbin, R. & Winn, J. A Bayesian framework to account for complex non-genetic factors in gene expression levels greatly increases power in eQTL studies. PLOS Computational Biology 6, e1000770 (2010).

31. Li, S. et al. Detecting and correcting systematic variation in large-scale RNA sequencing data. Nature biotechnology 32, 888–895 (2014).

32. Gagnon-Bartsch, J.A. & Speed, T.P. Using control genes to correct for unwanted variation in microarray data. Biostatistics 13, 539–552 (2012).

33. Risso, D., Schwartz, K., Sherlock, G. & Dudoit, S. GC-content normalization for RNA-Seq data. BMC Bioinformatics 12, 1–17 (2011).

34. Hansen, K.D., Irizarry, R.A. & Wu, Z. Removing technical variability in RNA-seq data using conditional quantile normalization. Biostatistics 13, 204–216 (2012).

35. Leek, J.T. & Storey, J.D. Capturing Heterogeneity in Gene Expression Studies by Surrogate Variable Analysis. PLOS Genetics 3, e161 (2007).

36. Leek, J.T., Johnson, W.E., Parker, H.S., Jaffe, A.E. & Storey, J.D. The sva package for removing batch effects and other unwanted variation in high-throughput experiments. Bioinformatics 28, 882–883 (2012).

37. Leek, J.T. Svaseq: removing batch effects and other unwanted noise from sequencing data. Nucleic acids research 42, e161–e161 (2014).

38. Stegle, O., Parts, L., Piipari, M., Winn, J. & Durbin, R. Using probabilistic estimation of expression residuals (PEER) to obtain increased power and interpretability of gene expression analyses. Nature protocols 7, 500–507 (2012).

39. Molania, R., Gagnon-Bartsch, J.A., Dobrovic, A. & Speed, T.P. A new normalization for Nanostring nCounter gene expression data. Nucleic acids research 47, 6073–6083 (2019).

40. Gagnon-Bartsch, J.A., Jacob, L. & Speed, T.P. Removing unwanted variation from high dimensional data with negative controls. Berkeley: Tech Reports from Dep Stat Univ California, 1–112 (2013).

41. Risso, D., Ngai, J., Speed, T.P. & Dudoit, S. Normalization of RNA-seq data using factor analysis of control genes or samples. Nature biotechnology 32, 896–902 (2014).

42. Dahl, A., Guillemot, V., Mefford, J., Aschard, H. & Zaitlen, N. Adjusting for principal components of molecular phenotypes induces replicating false positives. Genetics 211, 1179–1189 (2019).

43. Jaffe, A.E. et al. Practical impacts of genomic data “cleaning” on biological discovery using surrogate variable analysis. BMC Bioinformatics 16, 1–10 (2015).

44. Gandal, M.J. et al. Transcriptome-wide isoform-level dysregulation in ASD, schizophrenia, and bipolar disorder. Science 362, eaat8127 (2018).

45. Carrico, A.W. et al. Recent stimulant use and leukocyte gene expression in methamphetamine users with treated HIV infection. Brain, behavior, and immunity 71, 108–115 (2018).

46. Fass, S.B. et al. Relationship between sex biases in gene expression and sex biases in autism and Alzheimer’s disease. Biology of Sex Differences 15, 47 (2024).

47. Van Rheenen, W. et al. Whole blood transcriptome analysis in amyotrophic lateral sclerosis: A biomarker study. PLOS ONE 13, e0198874 (2018).

48. Parikshak, N.N. et al. Genome-wide changes in lncRNA, splicing, and regional gene expression patterns in autism. Nature 540, 423–427 (2016).

49. Saitou, M., Dahl, A., Wang, Q. & Liu, X. Allele frequency impacts the cross-ancestry portability of gene expression prediction in lymphoblastoid cell lines. The American Journal of Human Genetics 111, 2814–2825 (2024).

50. Lee, M. et al. Circulating MicroRNA expression levels associated with internet gaming disorder. Frontiers in Psychiatry 9, 81 (2018).

51. Gross, T.J. et al. Plasma metabolites related to cellular energy metabolism are altered in adults with Down syndrome and Alzheimer’s disease. Developmental Neurobiology 79, 622–638 (2019).

52. Ellis, S.E., Panitch, R., West, A.B. & Arking, D.E. Transcriptome analysis of cortical tissue reveals shared sets of downregulated genes in autism and schizophrenia. Transl Psychiatry 6, e817–e817 (2016).

53. Voisey, J. et al. Clinically proven drug targets differentially expressed in the prefrontal cortex of schizophrenia patients. Brain, behavior, and immunity 61, 259–265 (2017).

54. Lane, J. et al. Impact of statins on gene expression in human lung tissues. PLOS ONE 10, e0142037 (2015).

55. Kriegeskorte, N., Simmons, W.K., Bellgowan, P.S. & Baker, C.I. Circular analysis in systems neuroscience: the dangers of double dipping. Nature neuroscience 12, 535–540 (2009).

56. Hoffman, G.E. & Schadt, E.E. variancePartition: interpreting drivers of variation in complex gene expression studies. BMC Bioinformatics 17, 1–13 (2016).

57. Buja, A. & Eyuboglu, N. Remarks on Parallel Analysis. Multivariate Behavioral Research 27, 509–540 (1992).

58. Leek, J.T. Asymptotic Conditional Singular Value Decomposition for High-Dimensional Genomic Data. Biometrics 67, 344–352 (2011).

59. Leek, J.T. & Storey, J.D. A general framework for multiple testing dependence. Proceedings of the National Academy of Sciences 105, 18718–18723 (2008).

60. Molania, R. et al. Removing unwanted variation from large-scale RNA sequencing data with PRPS. Nature Biotechnology 41, 82–95 (2023).

61. Storey, J.D. A direct approach to false discovery rates. Journal of the Royal Statistical Society Series B: Statistical Methodology 64, 479–498 (2002).

62. A. Bennett, D.A. Schneider, J., Arvanitakis, Z. & S. Wilson, R. Overview and Findings from the Religious Orders Study. Current Alzheimer Research 9, 628–645 (2012).

63. A. Bennett, D. et al. Overview and Findings from the Rush Memory and Aging Project. Current Alzheimer Research 9, 646–663 (2012).

64. Wang, M. et al. The Mount Sinai cohort of large-scale genomic, transcriptomic and proteomic data in Alzheimer’s disease. Sci Data 5, 180185 (2018).

65. Hoffman, G.E. et al. CommonMind Consortium provides transcriptomic and epigenomic data for Schizophrenia and Bipolar Disorder. Sci Data 6, 180 (2019).

66. Fromer, M. et al. Gene expression elucidates functional impact of polygenic risk for schizophrenia. Nature neuroscience 19, 1442–1453 (2016).

67. Duke, D., Moran, L., Pearce, R. & Graeber, M. The medial and lateral substantia nigra in Parkinson’s disease: mRNA profiles associated with higher brain tissue vulnerability. Neurogenetics 8, 83–94 (2007).

68. Lesnick, T.G. et al. A genomic pathway approach to a complex disease: axon guidance and Parkinson disease. PLOS Genetics 3, e98 (2007).

69. Zheng, B. et al. PGC-1 α, a potential therapeutic target for early intervention in Parkinson’s disease. Science translational medicine 2, 52ra73–52ra73 (2010).

70. Zhang, Y., James, M., Middleton, F.A. & Davis, R.L. Transcriptional analysis of multiple brain regions in Parkinson’s disease supports the involvement of specific protein processing, energy metabolism, and signaling pathways, and suggests novel disease mechanisms. American Journal of Medical Genetics Part B: Neuropsychiatric Genetics 137, 5–16 (2005).

71. Dijkstra, A.A. et al. Evidence for immune response, axonal dysfunction and reduced endocytosis in the substantia nigra in early stage Parkinson’s disease. PLOS ONE 10, e0128651 (2015).

72. Ganna, A., Lee, D., Ingelsson, E. & Pawitan, Y. Rediscovery rate estimation for assessing the validation of significant findings in high-throughput studies. Briefings in Bioinformatics 16, 563–575 (2015).

73. Chen, J. et al. Fast and robust adjustment of cell mixtures in epigenome-wide association studies with SmartSVA. BMC Genomics 18, 1–13 (2017).

74. Zhang, B. et al. Integrated systems approach identifies genetic nodes and networks in late-onset Alzheimer’s disease. Cell 153, 707–720 (2013).

75. Satoh, J.-i., Yamamoto, Y., Asahina, N., Kitano, S. & Kino, Y. RNALSeq Data Mining: Downregulation of NeuroD6 Serves as a Possible Biomarker for Alzheimer’s Disease Brains. Disease markers 2014, 123165 (2014).

76. Miller, J.A., Woltjer, R.L., Goodenbour, J.M., Horvath, S. & Geschwind, D.H. Genes and pathways underlying regional and cell type changes in Alzheimer’s disease. Genome medicine 5, 48 (2013).

77. Avramopoulos, D., Szymanski, M., Wang, R. & Bassett, S. Gene expression reveals overlap between normal aging and Alzheimer’s disease genes. Neurobiol Aging 32, 2319. e2327-2319. e2334 (2011).

78. Blalock, E.M. et al. Incipient Alzheimer’s disease: microarray correlation analyses reveal major transcriptional and tumor suppressor responses. Proceedings of the National Academy of Sciences 101, 2173–2178 (2004).

79. Colangelo, V. et al. Gene expression profiling of 12633 genes in Alzheimer hippocampal CA1: transcription and neurotrophic factor downLregulation and upLregulation of apoptotic and proLinflammatory signaling. Journal of neuroscience research 70, 462–473 (2002).

80. Liang, W.S. et al. Altered neuronal gene expression in brain regions differentially affected by Alzheimer’s disease: a reference data set. Physiological genomics 33, 240–256 (2008).

81. Mostafavi, S. et al. A molecular network of the aging human brain provides insights into the pathology and cognitive decline of Alzheimer’s disease. Nature neuroscience 21, 811–819 (2018).

82. Allen, M. et al. Conserved brain myelination networks are altered in Alzheimer’s and other neurodegenerative diseases. Alzheimer’s & dementia 14, 352–366 (2018).

83. Choi, K.H. et al. Gene expression and genetic variation data implicate PCLO in bipolar disorder. Biological psychiatry 69, 353–359 (2011).

84. Chu, T.T., Liu, Y. & Kemether, E. Thalamic transcriptome screening in three psychiatric states. Journal of human genetics 54, 665–675 (2009).

85. Padmos, R.C. et al. A discriminating messenger RNA signature for bipolar disorder formed by an aberrant expression of inflammatory genes in monocytes. Archives of General Psychiatry 65, 395–407 (2008).

86. Ryan, M. et al. Gene expression analysis of bipolar disorder reveals downregulation of the ubiquitin cycle and alterations in synaptic genes. Molecular psychiatry 11, 965–978 (2006).

87. Wu, J.Q. et al. Transcriptome sequencing revealed significant alteration of cortical promoter usage and splicing in schizophrenia. PLOS ONE 7, e36351 (2012).

88. Hwang, Y. et al. Gene expression profiling by mRNA sequencing reveals increased expression of immune/inflammation-related genes in the hippocampus of individuals with schizophrenia. Transl Psychiatry 3, e321–e321 (2013).

89. Collado-Torres, L. et al. Regional heterogeneity in gene expression, regulation, and coherence in the frontal cortex and hippocampus across development and schizophrenia. Neuron 103, 203-216. e208 (2019).

90. Manchia, M. et al. Pattern of gene expression in different stages of schizophrenia: Down-regulation of NPTX2 gene revealed by a meta-analysis of microarray datasets. European Neuropsychopharmacology 27, 1054–1063 (2017).

91. Parker, H.S. et al. Preserving biological heterogeneity with a permuted surrogate variable analysis for genomics batch correction. Bioinformatics 30, 2757–2763 (2014).

92. Chakraborty, S., Datta, S. & Datta, S. svapls: an R package to correct for hidden factors of variability in gene expression studies. BMC Bioinformatics 14, 236 (2013).

93. Lee, S., Sun, W., Wright, F.A. & Zou, F. An improved and explicit surrogate variable analysis procedure by coefficient adjustment. Biometrika 104, 303–316 (2017).

94. Siska, C. & Kechris, K. Differential correlation for sequencing data. BMC research notes 10, 54 (2017).

95. Leek, J.T. et al. Package ‘sva’. (2014).

96. Ritchie, M.E. et al. limma powers differential expression analyses for RNA-sequencing and microarray studies. Nucleic acids research 43, e47–e47 (2015).

97. Benjamini, Y. & Hochberg, Y. Controlling the false discovery rate: a practical and powerful approach to multiple testing. Journal of the Royal statistical society: series B (Methodological) 57, 289–300 (1995).

98. Baratloo, A., Hosseini, M., Negida, A. & El Ashal, G. Part 1: simple definition and calculation of accuracy, sensitivity and specificity. (2015).

99. Molinaro, A.M. Diagnostic tests: how to estimate the positive predictive value. Neuro-Oncology Practice 2, 162–166 (2015).

100. A Bennett, D. et al. Overview and findings from the rush Memory and Aging Project. Current Alzheimer Research 9, 646–663 (2012).

101. Wang, M. et al. The Mount Sinai cohort of large-scale genomic, transcriptomic and proteomic data in Alzheimer’s disease. Sci Data 5, 1–16 (2018).

102. Hoffman, G.E. et al. CommonMind Consortium provides transcriptomic and epigenomic data for Schizophrenia and Bipolar Disorder. Sci Data 6, 180 (2019).

103. Moran, L. et al. Whole genome expression profiling of the medial and lateral substantia nigra in Parkinson’s disease. Neurogenetics 7, 1–11 (2006).

104. Wang, Q. et al. The landscape of multiscale transcriptomic networks and key regulators in Parkinson’s disease. Nature Communications 10, 5234 (2019).

105. Robinson, M.D., McCarthy, D.J. & Smyth, G.K. edgeR: a Bioconductor package for differential expression analysis of digital gene expression data. Bioinformatics 26, 139–140 (2010).

106. Wang, Q. et al. The landscape of multiscale transcriptomic networks and key regulators in Parkinson’s disease. Nature Communications 10 (2019).

107. Mirra, S.S. et al. The Consortium to Establish a Registry for Alzheimer’s Disease (CERAD) Part II. Standardization of the neuropathologic assessment of Alzheimer’s disease. Neurology 41, 479–479 (1991).

108. Han, S. et al. Whole-genome sequencing analysis of suicide deaths integrating brain-regulatory eQTLs data to identify risk loci and genes. Molecular psychiatry 28, 3909–3919 (2023).

109. Team, R.C. (R Foundation for Statistical Computing, Vienna, Austria; 2022).

110. Gentleman, R.C. et al. Bioconductor: open software development for computational biology and bioinformatics. Genome biology 5, 1–16 (2004).

111. Huber, W. et al. Orchestrating high-throughput genomic analysis with Bioconductor. Nat Methods 12, 115–121 (2015).

112. Microsoft; Weston, S., Edn. 1.5.2 (2022).

113. Durinck, S., Spellman, P.T., Birney, E. & Huber, W. Mapping identifiers for the integration of genomic datasets with the R/Bioconductor package biomaRt. Nature protocols 4, 1184–1191 (2009).

114. Davis, S. & Meltzer, P.S. GEOquery: a bridge between the Gene Expression Omnibus (GEO) and BioConductor. Bioinformatics 23, 1846–1847 (2007).

115. Wickham, H. et al. Welcome to the Tidyverse. Journal of open source software 4, 1686 (2019).

116. Dowle, M. et al. Package ‘data. table’. Extension of ‘data. frame 596, 952 (2019).

117. Wickham, H. et al. Package ‘readr’. Read Rectangular Text Data. Available online: https://cran.r-project.org/web/packages/readr/readr.pdf (accessed on 23 August 2023) (2024).

118. Wickham, H. & Wickham, H. Data analysis. (Springer, 2016).

119. Petukhov, V., van den Brand, T. & Biederstedt, E. (2020).

120. Slowikowski, K. et al. Package ggrepel. Automatically position non-overlapping text labels with ‘ggplot2 (2018).

121. Garnier, S. et al. Package ‘viridis’. Colorblind-Friendly Color Maps for R (2018).

122. Pedersen, T.L. Package ‘patchwork’. R package http://CRAN.R-project.org/package=patchwork.Cran (2019).

123. Borg, I. & Groenen, P.J. Modern multidimensional scaling: Theory and applications. (Springer, 2005).

124. Scott, D.W. (Taylor & Francis, 1990).

125. Hahsler, M., Hornik, K. & Buchta, C. Getting things in order: an introduction to the R package seriation. Journal of Statistical Software 25, 1–34 (2008).

