## Supplementary Information for "Surrogate variables and the risk of overfitting in neuropsychiatric gene expression analyses"

### SVA generally estimates the number of simulated EIVs accurately

To evaluate the accuracy of SVA methods in estimating the number of EIVs, the estimated number of SVs were compared to the known number of EIVs (***Extended Data Fig. 1A***). Both SVA methods generally performed well at estimating the true number of EIVs. The difference between the true number of EIVs and the median estimated number of SVs ranged from 0 to 22 for SVA “BE” and from 0 to 18 for SVA “Leek.” The “Leek” method underestimated the number of EIVs in 77.32% of simulations and correctly estimated the number in 22.68% of simulations (***Extended Data Fig. 1A***). The “BE” method, while generally underestimating the number of EIVs, correctly estimated the number in 8.17% of simulations, underestimated in 90.14%, and overestimated in 1.69% (***Extended Data Fig. 1A***). In the case where 0 EIVs were simulated, both SVA “BE” and “Leek” methods were able to accurately detect the presence of 0 EIVs in 99.36% and 100% of 9,047 simulations respectively (0.34% of all simulations). For the 58 instances where SVA “BE” incorrectly estimated presence of >0 EIVs, only one SV was estimated to be present.

The relationship between the number of EIVs and the FPR was also characterized (***Extended Data Fig. 1B***). Strikingly, while the median FPR of DE analyses on synthetic data without any EIVs was 0.004, with as low as 10 EIVs, the FPR drastically increased (median FPR = 48.3%, 48.1%, and 47.4% for “Known number of SVs,” SVA “BE,” and “Leek” respectively). FPR generally decreased as the number of EIVs increased to 100 (***Extended Data Fig. 1B***), consistent with a reduction in the number of total positives for the DEA (***EIV 10, median number of TPs= 1558.8, EIV 100, median number of TPs = 1519.9***), while the number of true positives remained comparable (***Supplementary Table 1)***. Furthermore, the number of true negatives generally increased, and the number of false positives generally decreased (***Supplementary Table 1***), leading to a reduction in FPR across all methods using SVs. Consistent with this, for all methods using SVs in DEA, FPRs generally decreased with an increasing number of EIVs for a given number of true simulated DEGs (***Supplementary Table 1***).

### Despite overfitting protection, SVs remain nearly orthogonal to phenotype

Given that the SVA procedure allows for SVs to be related to the phenotype of interest to prevent overfitting, it is essential to assess the degree of orthogonality of the angles of the SV vectors inferred from the data with knowledge of the phenotype of interest with the vector of the phenotype of interest. For this, the cosine angle between the SV vectors and the phenotype vector was calculated (See *Methods*). For every neuropsychiatric disorder dataset, the SV vectors were found to be approximately orthogonal to the phenotype of interest (***Extended Data Fig. 6A***). This result indicates that SVs inferred from real data are capturing little to no variation in the data explained by the phenotype of interest, potentially contributing to the overfitting observed when SVA is used in DEA.

SVA does not lead to overfit under null conditions

To assess whether overfitting is only observed when phenotype of the interest is present, we evaluated the impact of SVA on DEG distributions under null conditions in ROSMAP (See *Methods*, ***Extended Data Fig. 6B***). Specifically, we performed a null analysis using shuffled group labels in the ROSMAP dataset and evaluated DEA performance with and without SVA under these null conditions. Under the shuffled conditions, where true biological signal is absent, FPR was well-controlled with SVA application, indicating that overfitting does not occur in the absence of phenotypic signal. This result demonstrates that overfitting observed in SVA is only present when true biological differences between groups exist rather than being an inherent methodological artifact, suggesting that the overfitting mechanism is linked to the ability of SVA to inadvertently capture phenotype-related variation during the SV estimation process.

### Our synthetic data framework reproduces original SVA results

To evaluate SVA performance, the author of the original SVA manuscript simulated 1,000 genes from 20 samples and introduced known batch effects along with a subset of DEGs to test the ability of SVA to distinguish true biological signal from technical artifacts while controlling for FPR (<https://jtleek.com/svaseq/simulateData.html>)^1^. To assess whether our larger-scale simulation setup would yield similar results at the same set of parameters, we adapted Leek's simulation parameters to match our study design. We scaled up from the original 1,000 genes and 20 samples to 20,000 genes and 1,000 samples while maintaining the exact proportions from the original simulation for every other parameter. For example, where Leek’s simulation included 200 DEGs and 800 non-DEGs with one SV affecting 40% of the genes, our scaled version included 4,000 DEGs and 16,000 non-DEGs with one SV affecting 40% of the genes. Using these scaled parameters, we found that SVA was able to maintain low. At parameters from the original SVA manuscript (total_eiv_sd = 2, status_sd = 1, resid_sd = 1) that matched our simulations (total_eiv_var = 1, status_var = 0.25, resid_var = 0.25), we observed that SVA maintained low FPR. Altogether, these results confirm that SVA maintains low FPR under conditions of minimal unwanted variation effects.

Convergence-based SV estimation reduces reproducibility of AD DE signatures

Following the development of SVA, an enhanced method called SmartSVA^2^ was introduced to address the false positive behaviors observed with traditional SVA. SmartSVA posits that SVA could lead to false positive due to its failure to reach convergence of SV estimation using a fixed number of iterations in its algorithm. SmartSVA fixes this by explicitly imposing a convergence criterion, in addition to softening the initial probability estimates of being affected by the primary variable to stabilize estimation under strong confounding and improving computational efficiency^2^.

To evaluate whether the dynamic convergence criteria of SmartSVA could reduce false positives in DEA, we tested how it affected DE signature reproducibility in AD using two real datasets (ROSMAP and MSBB). Like our previous approach with traditional SVA, we assessed reproducibility by calculating Spearman correlations of log_2_FC of DEGs between the two datasets. We compared three scenarios: no SVA correction, SmartSVA with “BE” method^3^, and SmartSVA with Random Matrix Theory (RMT) method^4, 5^. For each SmartSVA approach, we evaluated performance using either all selected SVs at once or by incrementally adding SV one at a time until reaching the maximum number selected by each method. We found that SmartSVA substantially reduced reproducibility compared to no SVA correction. The correlation dropped from r=0.46 (p < 2.2e^-16^) without SVA correction to r=0.12 (p < 2.2e^-16^) with “BE” and r=0.09 (p < 2.2e^-16^) with RMT when using all selected SVs. This reduced reproducibility was consistently observed even when adding SV incrementally one at a time (***Extended Data Figure 8***). The “BE” method selected 27 SVs for MSBB and 47 SVs for ROSMAP (***Extended Data Figure 8A***), while RMT selected 32 SVs for MSBB and 68 SVs for ROSMAP (***Extended Data Figure 8B***).

Notably, when using SmartSVA with “BE” at the maximum convergence criterion of B=100, MSBB achieved convergence while ROSMAP could not (the algorithm looped indefinitely), suggesting that convergence behavior might depends on dataset-specific characteristics, warranting further examination.

References

1.	Leek, J.T. & Storey, J.D. A general framework for multiple testing dependence. *Proceedings of the National Academy of Sciences* **105**, 18718-18723 (2008).

2.	Chen, J. et al. Fast and robust adjustment of cell mixtures in epigenome-wide association studies with SmartSVA. *BMC Genomics* **18**, 1-13 (2017).

3.	Buja, A. & Eyuboglu, N. Remarks on Parallel Analysis. *Multivariate Behavioral Research* **27**, 509-540 (1992).

4.	Plerou, V. et al. Random matrix approach to cross correlations in financial data. *Physical Review E* **65**, 066126 (2002).

5.	Teschendorff, A.E., Zhuang, J. & Widschwendter, M. Independent surrogate variable analysis to deconvolve confounding factors in large-scale microarray profiling studies. *Bioinformatics* **27**, 1496-1505 (2011).
